# Evidence for strong interplay between the nucleotide and base excision repair pathways in *D. radiodurans*

**DOI:** 10.64898/2026.06.04.729788

**Authors:** Mohammad Rida Hayek, Salvatore De Bonis, Christine Saint-Pierre, Jean-Baptiste Reiser, Elin Moe, Jean-Luc Ravanat, Joanna Timmins

## Abstract

*Deinococcus radiodurans* harbors a largely classical bacterial DNA repair machinery yet displays exceptional resistance to ultra-violet and ionizing radiation. To investigate whether crosstalk between its DNA repair pathways contributes to this phenotype, we mapped putative interactions between the nucleotide excision repair (NER) and base excision repair (BER) pathways, which together are responsible for the removal of nucleobase lesions. Using a bacterial two-hybrid system, we identified multiple direct interactions between NER and BER proteins, notably involving the two UvrA variants, and validated these interactions *in vitro*. Furthermore, functional analyses revealed that NER interferes with the BER-mediated removal of oxidized guanines by the Fpg DNA glycosylase, likely through competition for DNA binding and sequestration of Fpg. Finally, UvrB and UvrC were found to further process the Fpg incision product in an ATP-dependent, UvrA1-independent manner. Together, these results demonstrate a multi-level crosstalk between NER and BER in *D. radiodurans*, which may contribute to its extraordinary DNA repair capacity. To our knowledge, this represents the first evidence of such a complex interplay in bacteria.

## INTRODUCTION

The integrity of genomic DNA is constantly challenged by exogenous sources, such as ultra-violet (UV) and ionizing radiation (IR) (1, 2) or endogenous factors, like replication errors and reactive oxygen species (ROS) (1, 3, 4). These threats lead to the accumulation of a variety of DNA lesions, including abasic sites, base modifications, mismatches, intra- and inter-strand crosslinks, bulky DNA adducts as well as single- and double-strand breaks (DSBs) (1, 3–6). If left unrepaired, these DNA lesions lead to mutations, genomic instability and even cell death (2, 4).

To preserve genomic integrity, cells have evolved complex DNA repair pathways (7, 8). Depending on the type of DNA damage, at least six major repair pathways function to safeguard the genome: direct reversal repair (DRR), base excision repair (BER), nucleotide excision repair (NER), mismatch repair (MMR), homologous recombination (HR), and non-homologous end joining (NHEJ) (1, 7, 9). While these pathways were historically considered to operate independently, growing evidence coming notably from the study of eukaryotic systems, suggests that they might be interconnected, forming a network of transiently interacting proteins that cooperate to maintain genomic integrity. For instance, the MMR complex, MSH2-MSH3, has been shown to facilitate EXO1 recruitment to DSBs in human cells to promote HR (10, 11). NER and BER, traditionally associated with the elimination of bulky adducts and oxidative DNA lesions, respectively, have also been found to cooperate in eukaryotes and in *E. coli* for the repair of oxidative DNA damage (12–25). In particular, UV-DDB and XPC from the NER pathway have been shown to act as early recognition factors in the repair of oxidized guanines (8-oxoG), aiding the BER OGG1 DNA glycosylase to locate its substrate, notably in poorly accessible regions, but also stimulating the turnover of the repair enzymes (12–16). Similarly, in *E. coli*, both the Fpg DNA glycosylase from the BER pathway and the UvrABC factors from the NER pathway have been shown to contribute to the repair of oxidative DNA lesions (24, 25). Conversely, BER factors, such as NEIL1 DNA glycosylase, have also been shown to play a role in the repair of NER substrates and in particular bulky DNA adducts, including DNA inter-strand crosslinks (26–30), further underscoring the complexity and functional overlap between BER and NER repair pathways.

While crosstalk between repair pathways, particularly between NER and BER, has been the subject of several reports, the mechanisms underlying this interplay has remained largely unexplored in bacteria. *Deinococcus radiodurans* is a Gram-positive bacterium, well-known for its remarkable resistance to ionizing radiation, UV light, desiccation, and oxidative stress (31–35). Its resilience has been attributed to its multiple antioxidant systems, which collectively protect its proteome against ROS-induced DNA damage, and its highly efficient DNA repair machinery (33, 35–37). Indeed, despite possessing a relatively conserved and ‘classical’ DNA repair repertoire, *D. radiodurans* displays an extraordinary DNA repair capacity capable of repairing thousands of base modifications and UV-induced pyrimidine dimers, as well as hundreds of DSBs in just a few hours (35, 36, 38, 39). This suggests that the repair enzymes alone are probably not sufficient to explain this outstanding repair capacity, even though they have been reported to exhibit enhanced catalytic turnover rates and broader substrate specificity (35). Interestingly, a recent study has revealed the existence of a tight interaction network among DNA repair proteins involved in DSB repair in *D. radiodurans* (36), suggesting that its remarkable DNA repair efficiency may result from tightly coordinated DNA repair activities.

The genome of *D. radiodurans* encodes for a complete NER pathway composed of highly conserved UvrA1, UvrB, UvrC and UvrD factors (35, 40, 41), along with a second UvrA variant, UvrA2, which is missing the UvrB-binding domain found in canonical UvrA proteins, and whose function remains elusive (42). In addition, *D. radiodurans* possesses an expanded BER DNA glycosylase repertoire, including five uracil DNA glycosylases, two 3-methyladenine DNA glycosylases and four oxidative DNA damage glycosylases (31, 35). These enzymes are complemented by essential end-processing enzymes (Exonuclease III, DNA polymerase I and DNA ligase) to perform the final steps in the NER and BER pathways (31, 35). Using a bacterial two-hybrid system, we mapped the interactions between NER and BER factors and then confirmed the identified interactions *in vitro* using purified enzymes. Furthermore, we evaluated the putative interplay between the NER machinery and the Fpg DNA glycosylase for the repair of the most prevalent type of oxidized DNA damage, namely 8-oxoG. Altogether, these studies provide compelling evidence for several direct interactions between NER and BER factors as well as a functional interplay between these two repair pathways for the elimination of oxidized bases, with UvrA2 emerging as a key player in this process.

## MATERIAL AND METHODS

### Bacterial adenylate cyclase two-hybrid (BACTH) assay

Codon-optimized genes for expression in *E. coli* encoding all NER (UvrA1 [DR1771], UvrA2 [DRA0188], UvrB [DR2275], UvrC [DR1354], and UvrD [DR1775]) and BER factors (Mismatch specific uracil DNA glycosylase (MUG [DR0715]), Uracil-DNA N glycosylase (UNG [DR0689]), Thermophilic DNA glycosylase (ThDG [DR1751]), Hypothetic uracil DNA glycosylase (HypoUDG [DR0022]), Hypothetic uracil DNA glycosylase 2 (HypoUDG2 [DR1663]), 3-methyladenine DNA glycosylase II (AlkA1 [DR2074]), 3-methyladenine DNA glycosylase II (AlkA2 [DR2584]), Endonuclease III1 (EndoIII1 [DR2438]), Endonuclease III2 (EndoIII2 [DR0289]), Endonuclease III3 (EndoIII3 [DR0928]), Formamidopyrimidine DNA glycosylase (Fpg [DR0493]), A/G specific adenine DNA glycosylase (MutY [DR2285]), Exodeoxyribonuclease III (ExoIII [DR0354]), DNA polymerase I (DNA PolI [DR1707]) and the DNA ligase A (DNA Lig [DR2069]) were synthesized by Twist Biosciences, cloned into a standard shuttle vector and subsequently cloned into pEB355 and pEB354 plasmids to be expressed, with the exception of UvrA1, in fusion with the C-termini of the T18 and T25 domains, respectively, of adenylate cyclase (43). For UvrA1, the T18-UvrA1 and T25-UvrA1 constructs were not functional in the BACTH system, so instead UvrA1 was expressed in fusion with the N-termini of T18 and T25 (UvrA1-T18 and UvrA1-T25). All clones were verified by restriction digest and DNA sequencing (Genewiz/Azenta Life Sciences).

For the BACTH assay, the reporter strain BTH101 missing the gene coding for the endogenous adenylate cyclase (Cya^-^ strain) was co-transformed with various combinations of the pEB355 and pEB354 plasmids, the first encoding the T18-bait fusion protein and the second the T25-prey fusion protein. Empty plasmids expressing T18 or T25 alone were used as negative controls. Transformants were plated on LB-agar plates containing ampicillin (100 µg/mL) and kanamycin (50 µg/mL) and were incubated for 48h at 30°C, after which 5 colonies were picked and grown overnight at 30°C in a shaking incubator in 3 mL of LB medium supplemented with ampicillin, kanamycin and 0.5 mM IPTG to induce the expression. The next day, 3 µL from each culture were spotted in duplicate on lactose-containing MacConkey (Sigma) agar plates supplemented with ampicillin and kanamycin. Plates were incubated for 48h at 30°C. Lactose fermenting bacteria, resulting from bait-prey interaction, are red in color, while the absence of interaction results in beige colonies.

### Cloning, expression, and purification of the NER and BER proteins

Cloning, expression, and purification of *D. radiodurans* UvrA1, UvrB, UvrC (40, 44), UvrA2 (42), EndoIII (45, 46), Fpg (46), UNG (47) and DNA ligase A (48) have been described previously. All constructs were expressed in *E. coli* BL21 (DE3) cells as N-terminally His-tagged proteins and expression was typically induced at 20°C overnight or 37°C for 3-4 hours with 1 mM isopropyl-β-D-thiogalactopyranoside (IPTG). Briefly, all proteins were first purified on a nickel affinity column (HisTrap Fast Flow Cytiva or Ni-IDA from Macherey Nagel). This step was followed by a TEV cleavage at 4°C overnight to remove the His-tag, followed by a second nickel affinity column to eliminate the uncleaved protein and the His-tagged TEV protease. This was then followed by an ion exchange or heparin chromatography step to remove protein and nucleic acid contaminants and a size exclusion chromatography step to ensure the samples were homogeneous and free of any aggregation. To ensure stability of UvrA2, 0.5 mM ATP is maintained in all buffers (42). This is not the case for UvrA1.

### Pull-down assay

His-tagged-bait and non-tagged prey proteins were incubated together at a 1:5 ratio for 30 min on ice. This mixture was then diluted in Buffer 1 (75 mM KCl, 50 mM Tris-HCl pH8, 0.5 mM ATP, 1 mM MgCl_2_, 1 mM DTT) supplemented with 20 mM imidazole and then mixed with 100 µL of Ni-IDA (Macherey-Nagel) resin pre-equilibrated with Buffer 1. The mixture was then applied on a 700 µL spin-column and the flow-through was collected. The resin was then washed with 10 column volumes (CV) Buffer 1 supplemented with 20 mM imidazole and 10 CVs with Buffer 1 supplemented with 50 mM imidazole. The last CV (100 µL) of the 50 mM imidazole wash was collected separately for analysis and comparison, before elution with 6 CVs of Buffer 1 supplemented with 250 mM Imidazole. Elutions were collected in 3 separate fractions. To check for nonspecific binding of the non-tagged prey protein to the resin, we performed a control reaction in which the His-tagged bait protein was omitted. Similarly, we also performed control experiments with only the His-tagged bait protein. The presence of protein in the elution fractions was verified using Bradford reagent (Bio-Rad) and appropriate fractions were run on 10% TGX Stain-free polyacrylamide gels (Bio-Rad).

### Native gel electrophoresis

UvrA2 (bait), EndoIII1 (prey) and EndoIII2 (prey) proteins at a concentration of 50 µM were run on a 5% Tris-Borate-EDTA (TBE) pH8.3 native polyacrylamide gel alongside mixes of 50 µM UvrA2 with either 50 (1:1 ratio) or 200 µM (1:4 ratio) EndoIII1 or EndoIII2, incubated for 10 min prior to electrophoresis. Electrophoresis was conducted in 1XTBE buffer at 100 V for 1h at 4°C. To visualize the protein bands, the gel was stained with Instant Blue.

### Bio-Layer Interferometry (BLI)

Prior to BLI measurements, UvrA1 and UvrA2 were buffer exchanged into 30 mM Na-phosphate pH 8.0, 150 mM NaCl (for UvrA1) or 150 mM KCl (for UvrA2), 1 mM MgCl_2_, 1 mM DTT and 0.5 mM ATP (for UvrA2 only, to avoid protein precipitation), and biotinylated *in vitro* on ice for 2 h using a 20-fold molar excess of EZ-Link NHS-PEG_4_-Biotin (ThermoFisher Scientific). NHS-biotinylation was used because attempts to produce C-terminally biotinylated UvrA constructs (using Biotin Acceptor Peptide tag) were unsuccessful. The proteins were then buffer exchanged back into their respective buffers: 50 mM Tris-HCl pH 8.0, 150 mM NaCl, 2 mM MgCl_2_, 1 mM DTT and 5% glycerol for biotinylated UvrA1 and 50 mM Tris-HCl pH 8.0, 150 mM KCl, 1 mM MgCl_2_, 1 mM DTT, 0.1 mM EDTA, 0.5 mM ATP, 0.01% Brij 35 and 5% glycerol for biotinylated UvrA2. BLI analyses were conducted on an Octet RED 96e (Sartorius, former FortéBio) instrument using SA Biosensors (Sartorius) for ligand immobilization and 96-well microplates (Greiner) for sample and buffer deliveries. The total working volume for all samples and buffers was 0.2 mL per well, the shaking speed was fixed at 1000 rpm, and the plate temperature was set to 25°C. To minimize nonspecific binding, 0.02% Tween 20 was added to all buffer solutions and prior to each analysis, the biosensor tips were pre-wetted in ligand buffers. Immobilization of biotinylated UvrA1 or UvrA2 was performed after a baseline step of 120s, followed by ligand loading for 600s at a final sample concentration of 10 µg/mL to achieve a stable spectral shift of 1 nm. Kinetics analyses were then performed with an association step monitored for 600s with analyte samples serially diluted to concentrations ranging from 0 to 20 or 40 µM in Buffer A1 (50 mM Tris-HCl pH 8.0, 75 mM NaCl, 2 mM MgCl_2_, 1 mM DTT) for UvrA1 analytes or Buffer A2 (50 mM Tris-HCl pH 8.0, 75 mM KCl, 0.5 mM ATP, 1 mM MgCl_2_, 1 mM DTT) for UvrA2 analytes, followed by a dissociation step in the same analyte buffers for a further 900s. Reference biosensors without any bound ligand were processed in the same way to monitor nonspecific binding. All sensors were single-use. Binding sensorgrams were processed using Data Analysis HT 11.1. Signals from reference biosensors and zero concentration controls were subtracted from signals obtained for each functionalized biosensor and each analyte concentration. Association and dissociation curves were then extracted and a steady state analysis was performed. Individual experiments were fitted in GraphPad Prism 10 to a one-site specific binding model (with or without Hill slope) to determine the affinity constants.

### BER/NER incision assays

The DNA substrate (50mer-8oxodG) used for these assays was a 50 mer dsDNA with an internal 8-oxo-G modification incorporated in position 26 of the labeled strand, bearing a 5’ fluorescein moiety (6-FAM, green). The 8-oxoG containing oligonucleotide (5’-FAM-GACTACGTACTGTTACGGCTCCATC<u>X</u>CTACCGCAATCAGGCCAGATCTGC-3’) in which X=8-oxoG synthesized by Eurogentec and its complementary strand with a cytosine opposite the 8-oxoG synthesized by MWG Biotech were resuspended in 10 mM Tris-HCl pH 8.0, 50 mM NaCl and 0.5 mM EDTA at a concentration of 100 µM. The dsDNA substrate (at 50 µM) was prepared by annealing labeled and unlabeled complementary oligonucleotides at a 1:1.1 ratio, to have a slight excess of the unlabeled strand. The annealing reaction was heated to 98°C for 5 minutes and transferred directly into 1 L of water at 100°C, which was allowed to cool slowly at room temperature. A non-fluorescent 8-oxoG substrate was also prepared for MALDI-ToF Mass Spectrometry (MS) analysis using an unlabeled 8-oxoG 50mer DNA strand synthesized by Biomers. Gapped DNA (gDNA) mimicking the product of Fpg activity was prepared in a similar way by annealing two short oligonucleotides (MWG Biotech) corresponding respectively to the sequence on the 5’ (5’-GACTACGTACTGTTACGGCTCCATC-3’) and 3’ (5’-CTACCGCAATCAGGCCAGATCTGC-3’) sides of the 8-oxoG in the 50mer-8oxodG DNA, with the same 50mer complementary strand used above. The 5’ fragment included a terminal 3’ phosphate and the 3’ fragment a terminal 5’ phosphate. The 5’ fragment was ordered either with (for analysis on gel) or without (for MALDI-ToF MS analysis) a 5’ 6-FAM moiety (5-FAM-gDNA or gDNA). For incision assays, 25 nM DNA substrate was incubated with NER factors (25 nM UvrA1, 0.5 µM UvrB and 0.5 µM UvrC) and/or 25 nM Fpg at 37°C in 50 mM Tris-HCl pH7.5, 50 mM KCl, 5 mM DTT, 2.5 mM ATP and 2.5 mM MgCl_2_ supplemented with 2 μM BSA. NER activity was initiated by addition of the ATP, whereas BER activity was initiated by the addition of Fpg. The reaction volume was typically set to 100 μL, and at defined timepoints, reactions were stopped by mixing 10 μL of the reaction mix with 10 μL stop buffer (2x TBE, 8 M urea, 0.025% bromophenol blue, and 0.1% SDS) and subsequent heating of the samples to 95°C for 5 min. Reactions were then analyzed on 20% TBE-8M urea polyacrylamide gels prerun at 5 W/gel in 1xTBE buffer. The gels were run for 30 min and the DNA bands were visualized and quantified on a Chemidoc MP imager (Biorad) using the appropriate excitation light and detection filter for the green fluorophore. Each experiment was performed at least three times and the mean values and standard deviation were plotted using GraphPad Prism 10. Kinetics data of substrate processing were fitted to a two-phase exponential decay model in GraphPad Prism 10 (Y = Y_min_ + [(Y_0_-Y_min_) * Frac^Fast^] x exp^-kfast*X^ + [(Y_0_-Y_min_) * Frac^Slow^] * exp^-kslow*X^) corresponding to the sum of a fast and a slow exponential decay, where Y_0_ corresponds to the Y value at t=0, Y_min_ to the lower plateau, Frac^Fast^ and Frac^Slow^ to the fractions of fast and slow components of the reaction and k_fast_ and k_Slow_ to their respective rate constants. All fits were very good with R^2^ values above 0.95.

### MALDI-ToF mass spectrometry analysis

For MALDI-ToF MS analysis, incision reactions containing 2.5 pmol DNA in 100 µl reaction buffer were stopped at 60 min by heating samples to 95°C for 5 min and were desalted using Ziptip column with 0.2 µL C_18_ resin (Millipore). The columns were rinsed with 10 µL water to remove the reaction buffer. The DNA was eluted with 10 µL 50% acetonitrile. 1 μL of matrix solution 3-HPA mixed with 1 μL of oligonucleotide sample were spotted onto a polished stainless MALDI target plate (Bruker) and dried under vacuum. Mass spectra were obtained with a MALDI-ToF Microflex^TM^ spectrometer (Bruker) operated in negative ion mode.

### Fluorescence polarization assay

Equilibrium fluorescence polarization DNA binding assays were performed on a Clariostar (BMG Labtech) microplate reader, fitted with polarization filters. The DNA substrates consisted of 6-FAM 5’-labelled 50 mer dsDNA containing a fluorescein-conjugated thymine in position 26 (50mer-FdT), mimicking a NER substrate (40, 44), a 8-oxoG in position 26 (50mer-8oxodG) or intact DNA (50mer). 0 to 1 µM UvrA1 or 0 to 5 µM UvrA2 were titrated into 2 nM 6-FAM-labelled DNA in binding buffer composed of 50 mM Tris-HCl pH 7.5, 50 mM KCl, 5 mM DTT, 0.01% Tween 20, 2.5 mM MgCl_2_ supplemented with 0.1 mg/mL BSA. 2.5 mM ATP or AMP-PNP (Jena Biosciences) was also included in the reaction buffer. Reaction volumes were set to 50 µL. In all cases, after subtracting the polarization values obtained for DNA alone, the mean data from at least three independent experiments were fitted to a standard binding equation (Y = B_max_* X^h^/(K_d_^h^+X^h^)) assuming a single binding site with Hill slope (h) using GraphPad Prism 10, where B_max_ is the difference between the polarization of completely bound and completely free oligo, X is the protein concentration and K_d_ is the equilibrium dissociation constant. All fits were very good with R^2^ values above 0.98.

### AlphaFold3 prediction

Models of the different UvrA-BER complexes were predicted using the online AlphaFold3 server (https://alphafoldserver.com/) with 2 ATP ligands bound to each UvrA monomer (49, 50). For each prediction, 5 models were generated and ranked. The quality of the models was evaluated by the model confidence scores ipTM (interface predicted TM-score), pTM (predicted TM-score), pLDDT (predicted local distance difference test), and PAE (predicted aligned error). Based on these initial models, additional predictions of single domain interactions were subsequently run on the AlphaFold3 server using the insertion domain (ID) of either UvrA1 or UvrA2 together with the interacting domain from each of the BER partner proteins. All interaction interfaces were further analyzed using PDBePISA (51) and visualized in Chimera X (52). A detailed summary of the scores and characteristics of the predicted complexes is presented in supplementary Tables S1 and S2.

## RESULTS

### Mapping NER-BER interactions

We first screened for *in vivo* functional interactions between the five NER and fifteen BER factors using the bacterial two-hybrid (BACTH) assay (Fig. 1). For this purpose, an *E. coli* reporter strain was co-transformed with a plasmid encoding a NER factor fused to the adenylate cyclase T25 domain and a second plasmid encoding a BER factor fused to the adenylate cyclase T18 domain (Fig. 1A), and vice versa (Fig. 1B). In addition, the reporter strain was also co-transformed with each of the constructs in combination with empty T25 or T18 plasmids to eliminate false positives (supplementary Fig. S1). This allowed us to detect that, when fused with T18, HypoUDG systematically produced red colonies indicative of a positive interaction even when combined with an empty plasmid encoding T25 alone (Fig. 1A and Fig. S1). This construct was thus excluded from any further analysis. Several interactions involving the NER factors UvrA1 and UvrA2 were identified (Fig. 1C). UvrA2, in particular, was found to interact with five DNA glycosylases from the BER pathway, four of which are specialized in the repair of oxidized bases (Fpg, EndoIII1, EndoIII2 and EndoIII3) and one belonging to the Uracil DNA Glycosylase superfamily, UNG, specialized in the removal of uracil from DNA (Fig. 1A-B). Four out of five of these interactions were detected in both T18/T25 combinations, irrespective of the fused domain (T18 or T25). The UvrA2-EndoIII3 interaction, however, was only detected when UvrA2 was fused to T25 and EndoIII3 to T18. Similarly, UvrA1 was also found to interact with two BER factors, the Fpg DNA glycosylase and the end-processing DNA ligase A enzyme (Fig. 1). These interactions were detected in both T18/T25 combinations. The remaining Uvr proteins (UvrB, UvrC and UvrD) showed no interactions with the BER factors (Fig. 1).

**Figure 1.**
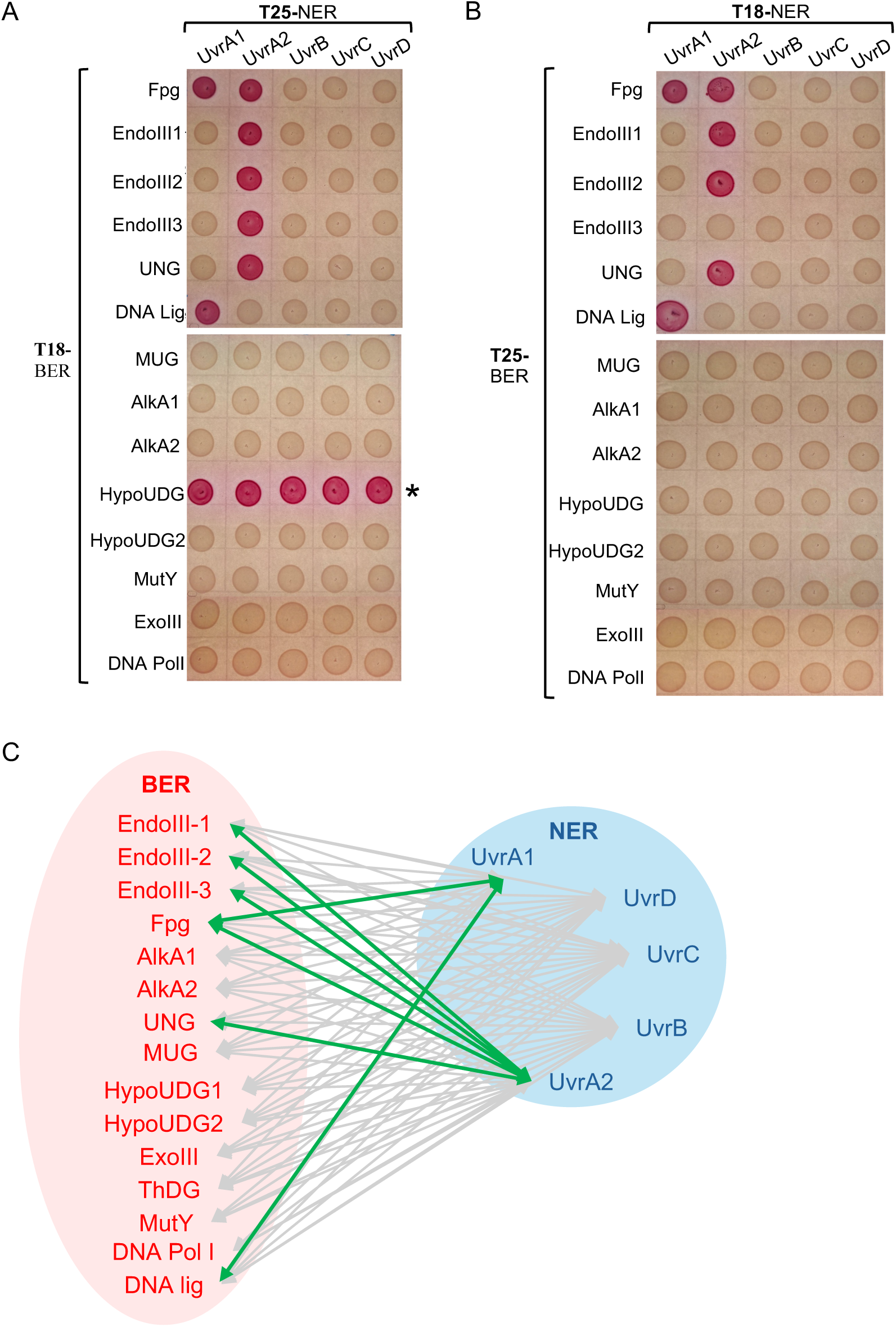
Bacterial two-hybrid analysis of the interactions between NER and BER factors. (A) Bait (BER) and prey (NER) factors fused respectively to the T18 and T25 domains of bacterial adenylate cyclase were co-transformed as indicated in the figure. (B) Bait (NER) and prey (BER) factors fused respectively to the T18 and T25 domains of bacterial adenylate cyclase were co-transformed as indicated in the figure. Positive interactions were detected on MacConkey agar plates as red-colored colonies. A false positive signal, indicated with a * was obtained with HypoUDG as a bait fused to T18. (C) Summary of the BER (red) - NER (blue) interactions investigated (grey lines) by BACTH in this study. The newly identified interactions that were subsequently validated *in vitro* are highlighted in green.

### *In vitro* validation of NER-BER interactions

To determine whether the interactions detected by BACTH were mediated by direct protein-protein interactions, we purified the relevant NER and BER factors and performed various biochemical and biophysical experiments. All seven NER-BER interactions were confirmed *in vitro* using at least one orthogonal approach. First, pull-down experiments were performed on all protein pairs (Fig. 2A-C, Fig. S2). Using His-tagged UvrA1, we confirmed the UvrA1-DNA ligase A interaction (Fig. 2A, Fig. S2A), but not the UvrA1-Fpg interaction (Fig. S2D). Purified DNA ligase, unlike Fpg, was indeed found to co-elute with His-tagged UvrA1. Next, using His-tagged UvrA2, we succeeded in validating the interactions of UvrA2 with EndoIII3 and UNG (Fig. 2B-C, Fig. S2B-C), both of which were found to co-elute with His-tagged UvrA2. In contrast, as for UvrA1, we were unable to detect the co-elution of His-tagged UvrA2 with Fpg (Fig. S2E). Moreover, the interaction of UvrA2 with either EndoIII1 or EndoIII2 could not be validated using this method due to non-specific binding of these BER enzymes to the nickel-affinity column (data not shown). This is likely due to the presence of a Fe-S cluster in these enzymes, which may favor their binding to metal-affinity resin. To overcome this issue, native gel electrophoresis was employed. For this purpose, UvrA2 alone or in the presence of an excess of EndoIII1 or EndoIII2 were separated by native gel electrophoresis (Fig. 2D-E). The UvrA2 band was partially shifted when incubated with a 5-fold excess of EndoIII1 and was fully shifted when incubated with a 20-fold excess of either EndoIII1 or EndoIII2. These results thus validate these interactions, but also indicate that they are likely weak.

**Figure 2.**
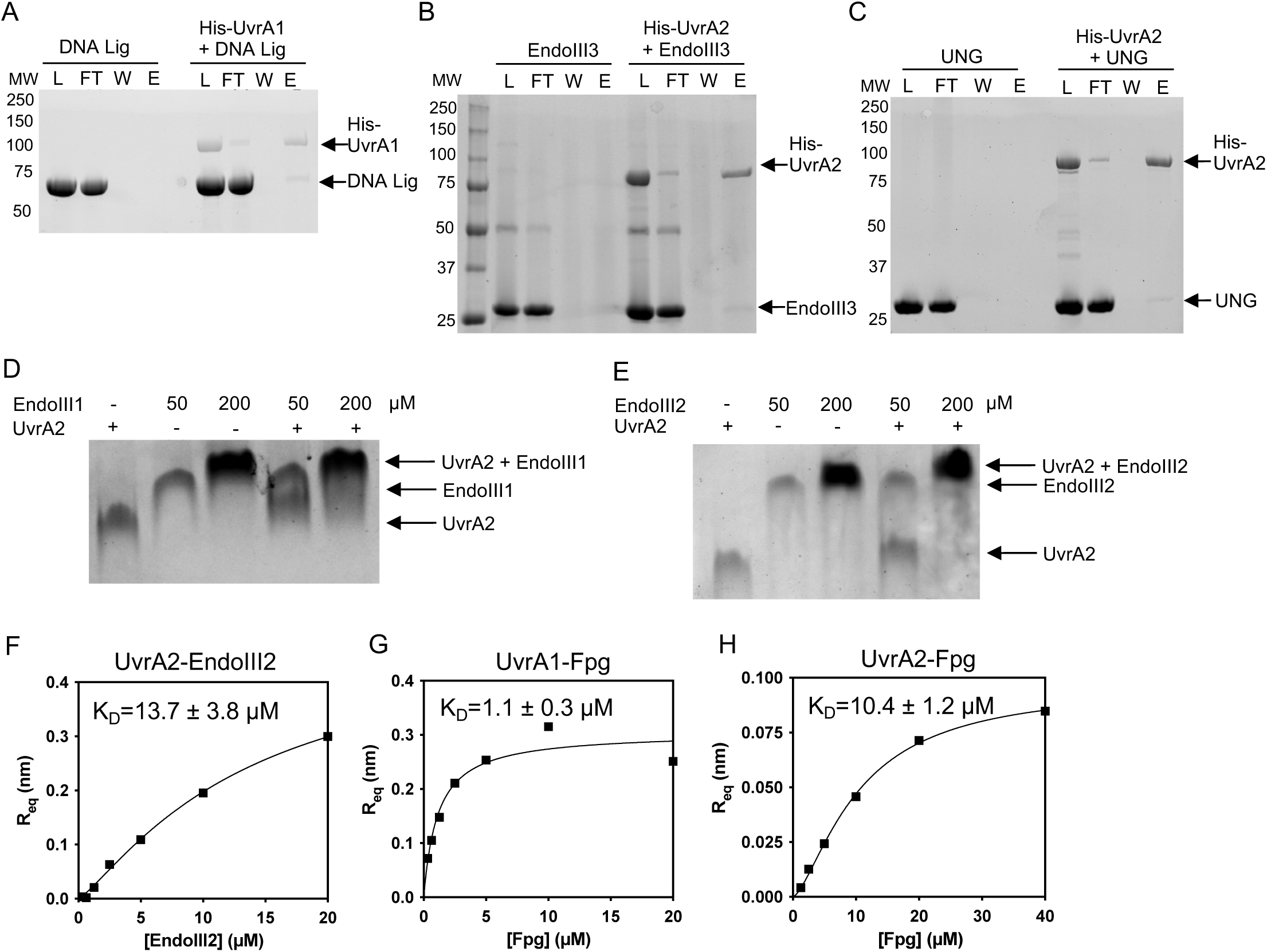
*In vitro* validation of identified NER-BER interactions. (A)-(C) SDS-PAGE analysis of the pull-down experiments validating the binding of His-tagged UvrA1 to DNA ligase (A) and of His-tagged-UvrA2 to EndoIII3 (B) and UNG (C). In each case, the first experiment (left) is a control experiment to verify that the non-tagged protein does not bind non-specifically to the Nickel affinity resin. Right: His-tagged UvrA1 or UvrA2 was mixed with a 5-fold molar excess of non-tagged DNA ligase, EndoIII3 or UNG and loaded (L) on Nickel affinity resin. The flow-through (FT) and the final wash (W) were collected before the protein was eluted (E). Molecular weight (MW) markers are indicated in kDa. (D)-(E) Electrophoretic Mobility Shift of UvrA2 binding to EndoIII1 (D) and EndoIII2 (E). Binding of EndoIII factors to UvrA2 cause the UvrA2 band to shift upwards in native TBE gel electrophoresis. Bands corresponding to the single and bound factors are indicated with arrows. (F)-(H) Steady-state analyses of the binding of EndoIII2 to UvrA2 (F) and of the binding of Fpg to immobilized UvrA1 (G) or UvrA2 (H) by BLI. Associated BLI sensorgrams are presented in Fig. S3. The presented data corresponds to one of two independent replicates. The data points were fitted to a one-site specific binding model with (for Fpg-UvrA binding) or without (UvrA2-EndoIII2) Hill slope in GraphPad Prism 10.

This was confirmed by bio-layer interferometry (BLI) experiments, which were performed first to evaluate the stability of UvrA2-EndoIII complexes and second, as described below, to assess the binding of the Fpg DNA glycosylase to the two UvrA factors, which we had failed to detect by either pull-down or native gel experiments (Fig. 2F-H and Fig. S3). For BLI measurements, biotinylated UvrA1 or UvrA2 was immobilized at the tips of streptavidin-coated sensors and BLI sensorgrams were subsequently recorded at a range of analyte concentrations (Fig. S3). We used BLI to compare the binding of the three EndoIII variants to UvrA2. Unfortunately, EndoIII1 exhibited non-specific binding to the empty sensor tip at high concentrations (*i.e.* 20 µM) and no binding to UvrA2 was detected at low concentrations (*i.e.* 1 µM), preventing any further analysis. Similarly, no interaction was detected for UvrA2-EndoIII3 even at 10 and 20 µM EndoIII3. For UvrA2-EndoIII2, however, we observed a concentration-dependent increase in the BLI response, confirming the interaction (Fig. 2F and Fig. S3A). Steady-state analysis was performed to evaluate the affinity of this interaction, which was estimated to be 13.7 ± 3.8 µM (Fig. 2F), confirming the transient nature of this interaction.

BLI was also performed to assess the binding of Fpg to both UvrA1 and UvrA2 (Fig. 2G-H). In both cases, we observed a concentration-dependent increase in the response signal, reflecting the specific binding of Fpg to biotinylated UvrA1 and UvrA2 (Fig. S3B-C), thereby validating our BACTH results. For UvrA1-Fpg, the BLI sensorgram was complex displaying an initial burst in the association signal, immediately followed by a rapid drop (Fig. S3B). This unusual profile was not suitable for kinetics analysis to determine association and dissociation constants. For UvrA2-Fpg, the binding profile was quite different, with a more conventional binding response (Fig. S3C). However, in this case, the association and dissociation kinetics were too rapid to extract reliable kinetic constants, so instead a steady state analysis was performed. These analyses yielded K_D_ values of 1.1 ± 0.3 µM for the UvrA1-Fpg interaction and of 10.4 ± 1.2 µM for UvrA2-Fpg. Fpg thus displays a significantly stronger binding to UvrA1 than to UvrA2.

In conclusion, all BER-NER interactions detected in BACTH were validated *in vitro* using purified proteins by at least one biochemical or biophysical method (Fig. 1C and Fig. 2), indicating that they rely on direct interactions between NER and BER factors, which appear to be weak and transient with K_D_ values in the micromolar range.

### Insertion domains of UvrA proteins are potential binding sites for DNA glycosylases

To gain insight into the molecular basis of the identified interactions, we employed AlphaFold3 to generate structural models of the UvrA (UvrA1 and UvrA2) dimers bound to monomeric BER enzymes (Fig. 3A-G). Although the predicted complexes exhibited low confidence scores (Table S1), a consistent binding pattern was observed across all the complexes. In a majority of the predicted complexes, the BER factors were indeed predicted to bind in between the flexible insertion domains (ID) of the two UvrA monomers and in close proximity to the ventral side of the core nucleotide-binding domains (NBD) of the UvrA dimers (Fig. 3 and Table S1). Interestingly, the confidence scores (ipTM) increased substantially when the predictions were re-iterated with only the interacting domains (Table S2). This was particularly striking for UvrA2-EndoIII models with ipTM values shifting from <0.2 to >0.5 and even >0.7 for UvrA2-EndoIII1 (Fig. 3H and Fig. S4). The predicted complexes exhibited large interaction surface areas involving numerous hydrogen bonds and salt bridges as revealed by PDBePISA analysis (Tables S1 and S2). The interacting area on each UvrA monomer was typically above 2000Å^2^ in the case of UvrA2 complexes, while it was lower (∼1000Å^2^) for UvrA1 complexes. In the case of UvrA2, the various DNA glycosylases were predicted to bind in a very similar way suggesting they may share a common binding site on UvrA2, and that binding is thus likely to be mutually exclusive (Fig. S4).

**Figure 3.**
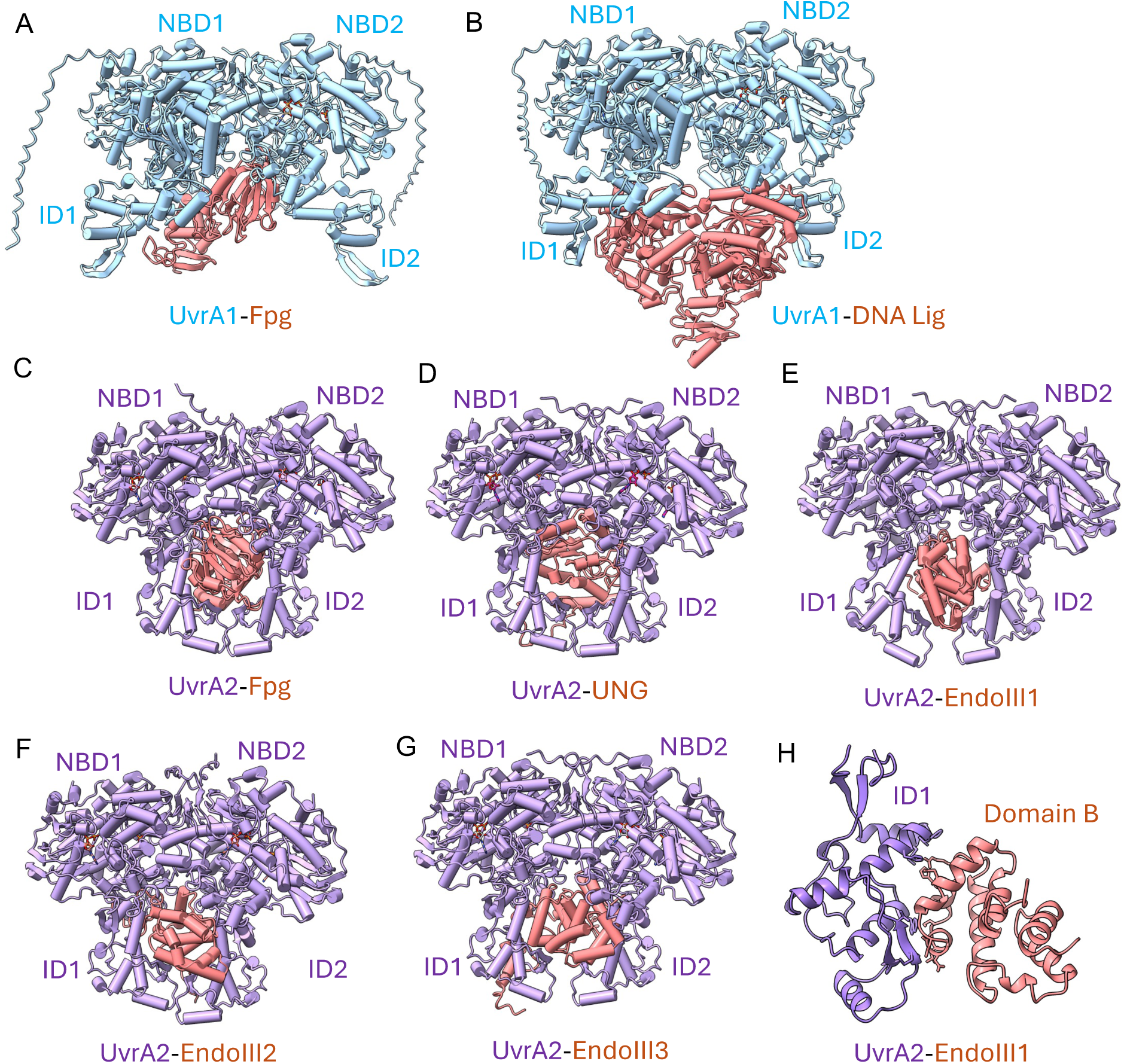
AlphaFold3 predicted models of UvrA dimers bound to BER factors. (A)-(B) Models of UvrA1 dimers (blue) bound to either Fpg (red, A) or DNA ligase (red, B). (C)-(G) Models of UvrA2 dimers (purple) bound to either Fpg (red, C), UNG (red, D), EndoIII1 (red, E), EndoIII2 (red, F) or EndoIII3 (red, G). The illustrated models correspond to the AF3 complexes exhibiting the highest interaction surface areas among the 5 models produced by AF3, and possessing the highest number of salt bridges and hydrogen bonds (Table S1). In all cases, the BER factors interact with the flexible insertion domains (ID) and the ventral side of the core nucleotide-binding domains (NBD) of the UvrA dimers. (H) AF3 Model of UvrA2 ID domain bound to domain B of EndoIII1.

### Functional interplay between NER and Fpg on oxidized DNA damage

We next wished to explore the potential functional interplay between the NER and BER pathways. For this, we focused on the repair of 8-oxoG, a highly mutagenic lesion that is also the most abundant form of oxidized base resulting from oxidative stress (53). When paired with a cytosine, 8-oxoG is very efficiently eliminated by the bifunctional DNA glycosylase Fpg. Fpg first hydrolyzes the glycosidic bond between the oxidized base and the ribose leading to the formation of an abasic site (AP), then cleaves the DNA backbone at this AP site via β- and δ-elimination, generating a single-strand break (1 nucleotide gap) with 3′- and 5′-phosphate termini. This cleavage reaction was followed by electrophoresis on denaturing urea gels (Fig. 4A) and the nature of the final products was confirmed by MALDI-ToF MS analysis (Fig. S5). When incubated alone with the 6-FAM 50 mer 8-oxoG substrate, Fpg processes 100% of the DNA within 5 min, releasing a 25 mer fragment, that is visible on the gel, corresponding to the DNA fragment released from the 5′ side of the 8-oxoG, and an unlabeled 24 mer fragment (detected only by MALDI-ToF, Fig. S5), corresponding to the DNA fragment released from the 3′ side of the 8-oxoG. Next, we treated this BER substrate with the minimal NER system, consisting of the UvrA1, UvrB and UvrC factors together with ATP (40). In this case, as expected, no incision was detected (Fig. 4B). Surprisingly, however, when the NER system was applied together with the Fpg DNA glycosylase, processing of the substrate by Fpg was significantly delayed and additional shorter DNA fragments were observed to accumulate ranging in size from ∼5 to 10 mer (Fig. 4C-D). The incision efficiency was reduced from 100% after 5 min to 75% after 1 h (Fig. 4D, Table 1), suggesting that the NER machinery may be competing with Fpg for binding to the oxidized DNA substrate, causing a marked delay in the incision of the DNA by Fpg.

**Figure 4.**
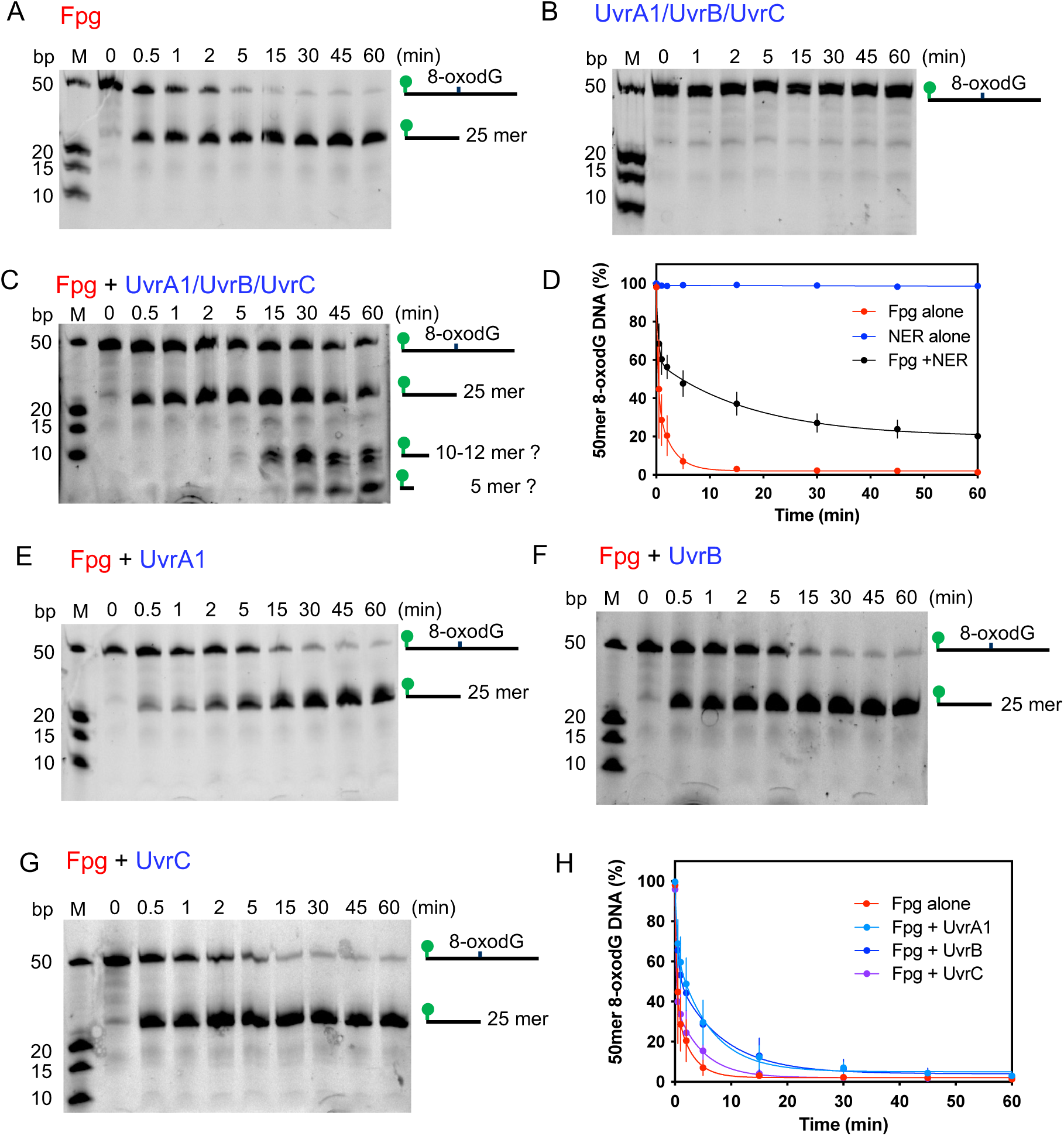
Synergistic activity of NER and Fpg on 8-oxoG-containing DNA substrate. (A)-(C) Representative urea polyacrylamide gels illustrating the incision activity of 25 nM Fpg (A), 25 nM UvrA1, 0.5 µM UvrB and 0.5 µM UvrC + 2.5 mM Mg^2+^/ATP (B) and 25 nM Fpg together with 25 nM UvrA1, 0.5 µM UvrB and 0.5 µM UvrC + 2.5 mM Mg^2+^/ATP (C) on 25 nM 50 mer 8-oxoG dsDNA. (A) Fpg activity leads to the accumulation of a 25 mer product bearing a 6-FAM moiety on its 5’ end. (B) The NER system is inactive on such a substrate. (C) The combined action of Fpg and the Uvr proteins causes a marked delay in the incision reaction and formation of the 25 mer fragment, and additional low molecular weight products ranging from 5 to 12 mer accumulated after 15 minutes. (D) Kinetics of incision of 8-oxoG DNA by either Fpg alone (red), NER alone (blue) or Fpg combined with the NER machinery (black). Data points corresponding to the mean of three independent measurements were fitted to a two-phase exponential decay model. Error bars correspond to the standard deviation. (E)-(G) Representative urea polyacrylamide gels illustrating the incision activity of 25 nM Fpg and 25 nM UvrA1 (E), 25 nM Fpg and 0.5 µM UvrB (F) and 25 nM Fpg and 0.5 µM UvrC (G). (H) Kinetics of incision of 8-oxoG DNA by either Fpg alone (red), Fpg + UvrA1 (light blue), Fpg + UvrB (dark blue) or Fpg + UvrC (purple). Data points corresponding to the mean of three independent measurements were fitted to a two-phase exponential decay model. Error bars correspond to the standard deviation.

**Table 1:** Rate constants of Fpg incision of 8-oxoG containing DNA in the absence and presence of the NER machinery.

| | Fixed $k_{Fast}$ ($\text{min}^{-1}$) | $k_{Slow}$ ($\text{min}^{-1}$) | % Fast/Slow | Plateau ($y_{min}$) % |
| --- | --- | --- | --- | --- |
| Fpg | $3.01 \pm 0.45$ | $0.36 \pm 0.15$ | 64/36 | $2.06 \pm 1.89$ |
| Fpg + UvrA1/UvrB/UvrC | $3.01 \pm 0.45$ | $0.06 \pm 0.02$ | 49/51 | $20.22 \pm 4.02$ |
| Fpg + UvrA1 | $3.01 \pm 0.45$ | $0.17 \pm 0.04$ | 35/65 | $5.11 \pm 2.13$ |
| Fpg + UvrB | $3.01 \pm 0.45$ | $0.15 \pm 0.04$ | 43/57 | $4.63 \pm 2.22$ |
| Fpg + UvrC | $3.01 \pm 0.45$ | $0.15 \pm 0.07$ | 68/32 | $1.64 \pm 2.21$ |
| Fpg + UvrB/UvrC | $3.01 \pm 0.45$ | $0.16 \pm 0.05$ | 54/46 | $4.48 \pm 2.17$ |
| Fpg + UvrB/UvrC (no ATP) | $3.01 \pm 0.45$ | $0.17 \pm 0.08$ | 60/40 | $29.20 \pm 2.13$ |
| Fpg + UvrA1 (AMP-PNP) | $3.01 \pm 0.45$ | $0.43 \pm 0.80$ | 81/19 | $3.88 \pm 2.32$ |
| Fpg + UvrA2 (ATP) | $3.01 \pm 0.45$ | $0.33 \pm 0.18$ | 69/31 | $3.15 \pm 1.91$ |
| Fpg + UvrA2 (AMP-PNP) | $3.01 \pm 0.45$ | ND | ND | ND |

For a more quantitative comparison, we fitted the kinetic measurements to a two-phase exponential decay model, in which the two components governing the rate of incision are: (i) the binding of Fpg to its substrate, and (ii) the incision reaction catalyzed by Fpg. To facilitate the comparison, based on prior knowledge (54), we made the assumption that once bound to its substrate, Fpg processes 8-oxoG at the same rate in all cases and we know that this activity is very efficient. Accordingly, we constrained the fast phase of the two-component decay (k_fast_) to a shared value in all fits (Table 1). We then assessed how the slower phase (k_Slow_, reflecting Fpg–substrate binding) and the residual amount of uncleaved DNA at reaction completion (the lower plateau, *y*ₘᵢₙ) were affected by adding NER factors to the reaction mixture. This analysis revealed that in the presence of the NER machinery, the k_Slow_ value was reduced 6-fold and that 20% of the initial substrate remained uncleaved after a 1h reaction (Table 1). Moreover, the appearance of additional lower molecular weight bands bearing the 5’ 6-FAM moiety, which were not observed when the NER and Fpg repair systems were applied separately, indicates that the NER machinery further processes the 5′ product of the Fpg incision, *i.e.* the 25 mer product with a 3′ phosphate.

### UvrC together with UvrB specifically incises the product of 8-oxoG elimination by Fpg

To further explore this synergistic NER-Fpg activity, we first performed a set of reactions in which each of the Uvr proteins were individually combined with Fpg (Fig. 4E-G). All three Uvr proteins caused a slight delay in the repair kinetics (k_Slow_ values were reduced two-fold), but, unlike the full NER system, all reactions finally reached completion after 1 hour, and no lower molecular weight bands were seen (Fig. 4H and Table 1). Next, we mixed Fpg with both UvrB and UvrC (Fig. 5A). Once again, we observed a slight delay in the incision activity of Fpg, similar to the effects of the individual UvrB and UvrC factors (Table 1). However, in addition, we also observed the appearance of low molecular weight DNA fragments (Fig. 5A), similar to that obtained with the complete NER machinery (Fig. 4C). This suggests that UvrB and UvrC together are responsible for the additional processing of the Fpg product, and that UvrA1 is dispensable for this activity. Interestingly, if ATP is left out of the reaction mix, the incision by Fpg is also delayed and almost 30% of the DNA substrate remains uncleaved at the end of the reaction, but the additional cleavage by UvrB/UvrC is not seen (Fig. 5B-C), suggesting that the weak ATP-dependent helicase activity of UvrB is needed for this cleavage reaction. The ATP may be needed to fuel the melting of the DNA duplex by the UvrB helicase, to provide UvrC with ssDNA substrate for its incision activity.

**Figure 5.**
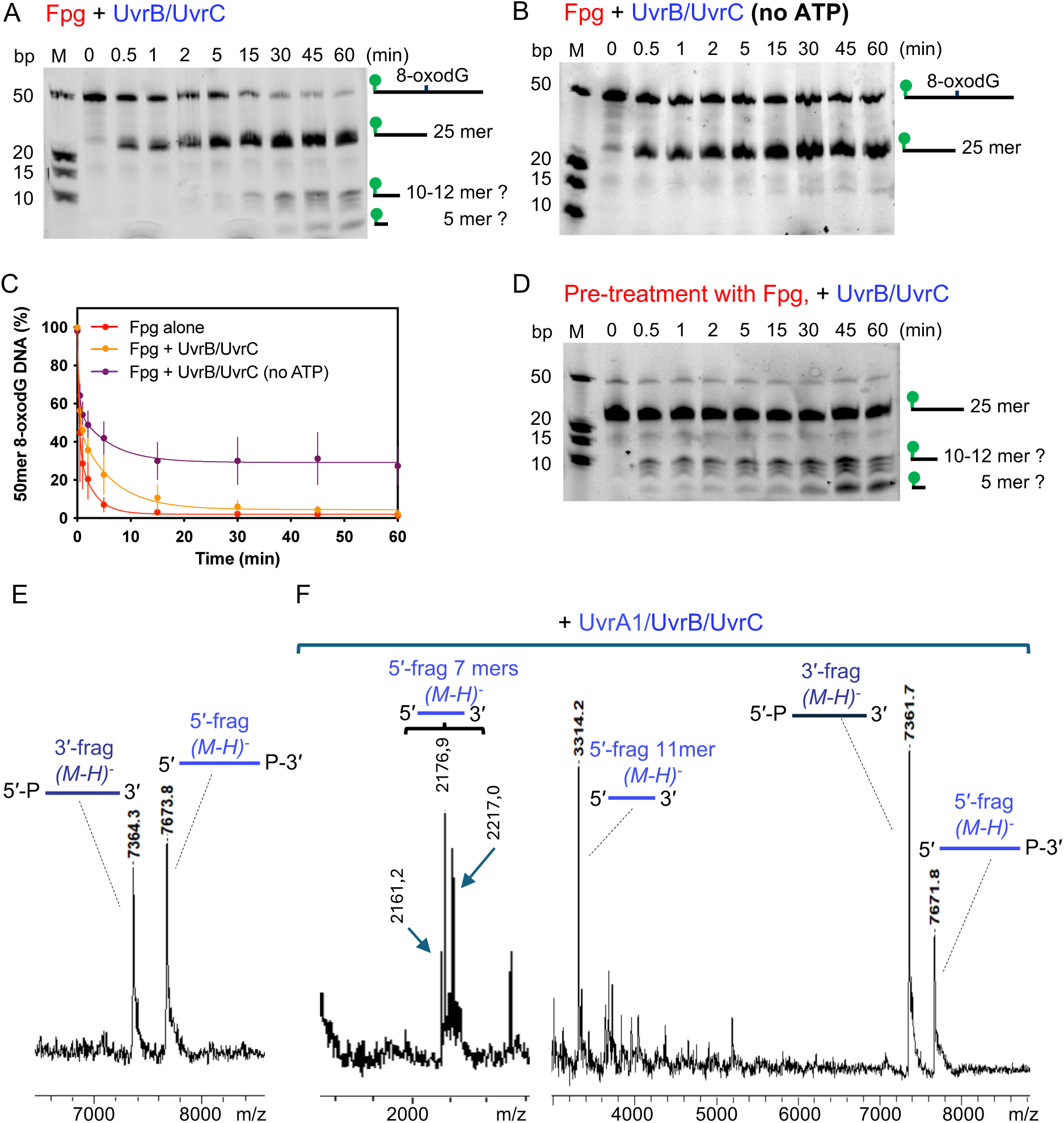
UvrB and UvrC processing of Fpg product. (A)-(B) Representative urea polyacrylamide gels illustrating the incision activity of 25 nM Fpg combined with 0.5 µM UvrB and 0.5 µM UvrC in the presence (A) and absence of ATP/MgCl_2_ (B). (C) Kinetics of incision of 8-oxoG DNA by either Fpg alone (red), Fpg + UvrB/UvrC + ATP (orange) or Fpg + UvrB/UvrC in the absence of ATP (purple). Data points corresponding to the mean of three independent measurements were fitted to a two-phase exponential decay model. Error bars correspond to the standard deviation. (D) Representative urea polyacrylamide gel illustrating the incision activity of 0.5 µM UvrB/UvrC on 8-oxoG DNA pre-treated with 25 nM Fpg for 20 min and heat-inactivated before addition of UvrB/UvrC and ATP to start the NER reaction. (E)-(F) MALDI-ToF MS spectra of the gapped DNA substrate (E) and the products of its processing by UvrA1/UvrB/UvrC in the presence of ATP (F). The nature of the main products is indicated (see also Table 2).

**Table 2:**
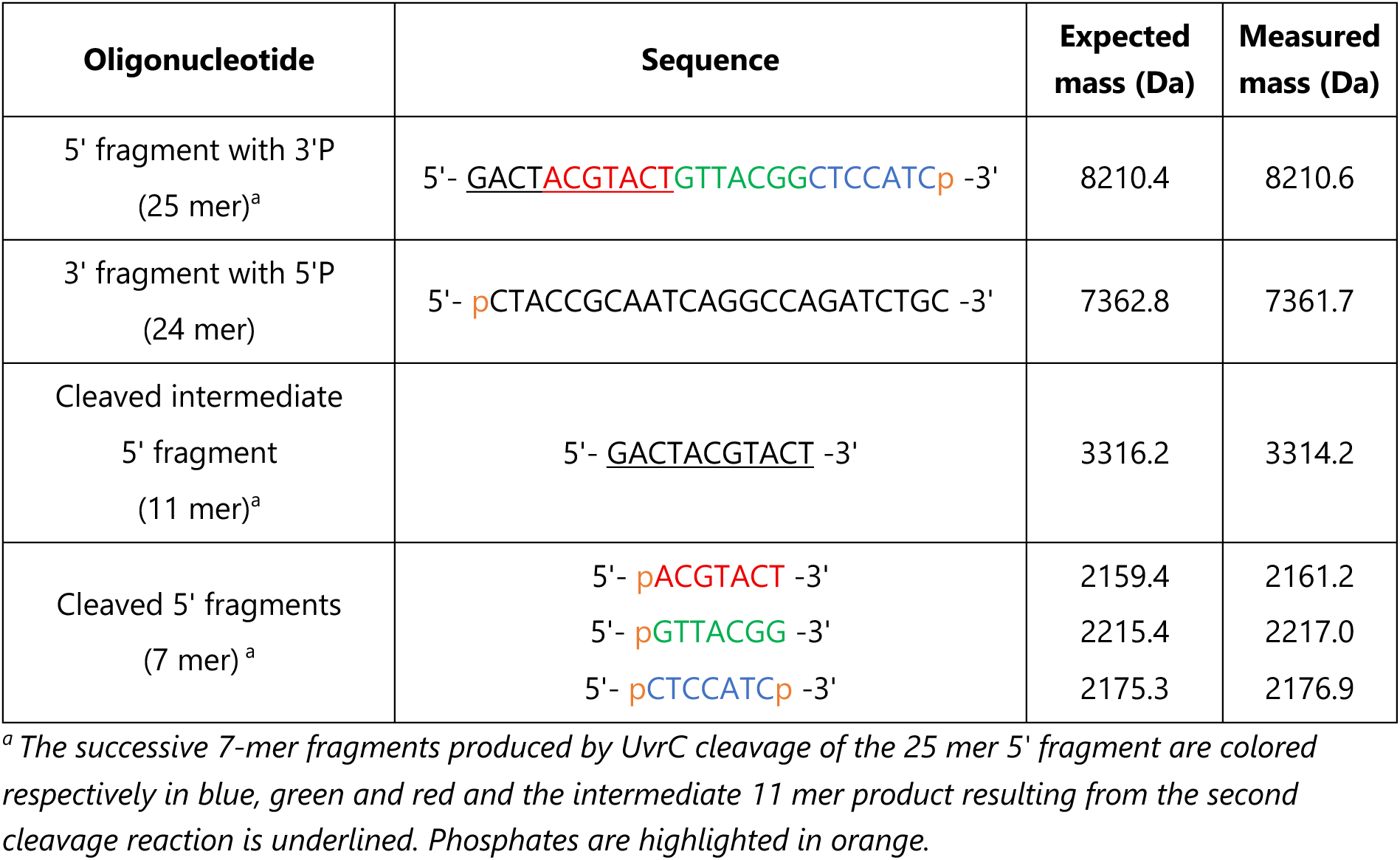
Expected and measured masses of DNA fragments determined by MALDI-ToF MS analysis after processing of 50 mer 1-nt gapped dsDNA by UvrB/UvrC.

Importantly, the UvrB/UvrC proteins were unable to process the 8-oxoG substrate in the absence of Fpg (Fig. S6A), indicating that DNA processing by Fpg is a prerequisite for further processing by the UvrB and UvrC enzymes. To verify this, we first treated the 8-oxoG substrate with Fpg, and then after heat inactivating the Fpg, we added the NER proteins. In these conditions, the lower molecular weight DNA fragments were found to rapidly accumulate, confirming our hypothesis that the Uvr proteins were actively processing the product of the Fpg incision (Fig. 5D). In addition, we saw the same cleavage pattern when incubating the NER machinery alone with a DNA substrate mimicking the product of the Fpg activity, with a one nucleotide (1-nt) gap in the position of the 8-oxoG (Fig. S5 and S6B). To determine the cleavage specificity, we analyzed the products of the reaction by MALDI-ToF MS. These analyses confirmed that the NER machinery specifically cleaves only the 25 mer fragment bearing a 3′ phosphate, positioned on the 5′ side of the 1-nt gap, irrespective of the presence or absence of a 6-FAM moiety on the 5′ extremity (Fig. 5E-F, Table 2). The 3′ fragment, in contrast, remained intact (Fig. 5E-F), as did the complementary strand. Interestingly, among the products, we identified three 7 mer products (bearing 5′ phosphate groups) and an intermediate 11 mer product corresponding to the products of successive cleavage reactions of the 25 mer 5′ fragment 7 nucleotides upstream of the gap (Fig. 5F, Table 2). To determine which of the two endonuclease domains of UvrC was responsible for this incision activity, experiments on gapped dsDNA were repeated with two UvrC point mutants, UvrC^E72A^ or UvrC^D391A^, that have previously been reported to be respectively impaired for either 3′ or 5′ cleavage (44). With UvrC^E72A^, the NER machinery efficiently cleaved the gapped DNA producing the same fragments as obtained with wild-type UvrC, as confirmed by MALDI-ToF analysis, while no cleavage was observed when UvrC^D391A^ was used (Fig. S6C). This clearly demonstrates that the C-terminal RNase H domain of UvrC is responsible for the processing of the Fpg gapped DNA product. Accordingly, this endonuclease domain is known to cleave 7 nucleotides upstream of DNA damage sites (44).

### Modulation of Fpg incision activity by UvrA1 and UvrA2 depends on their nucleotide-bound state and not their ability to recognize oxidative DNA damage

UvrA1, when added at equimolar concentration, caused a slight delay in the repair kinetics by Fpg (Fig. 4E and H, Table 1), suggesting that UvrA1 may compete for the binding to the 8-oxoG substrate. We repeated the experiment with UvrA2 instead of UvrA1, and found that UvrA2 did not cause any delay in the Fpg activity (Fig. 6A, Table 1). In earlier studies, the binding of UvrA1 and UvrA2 to damaged DNA has been shown to be differentially modulated by the nature of the nucleotide present in the reaction mix. UvrA1 differentiates between damaged and undamaged DNA more efficiently in the presence of ATP (40), whereas UvrA2 binds more tightly to damaged DNA in the presence of a non-hydrolyzable nucleotide, notably AMP-PNP (42). We, therefore, repeated the incision experiments with Fpg and UvrA1 or UvrA2 in the presence of AMP-PNP instead of ATP (Fig. 6B-C). Two strikingly different profiles were obtained. In the presence of AMP-PNP, Fpg incision was unaffected by UvrA1 and instead completely inhibited by UvrA2 (Fig. 6D, Table 1).

**Figure 6.**
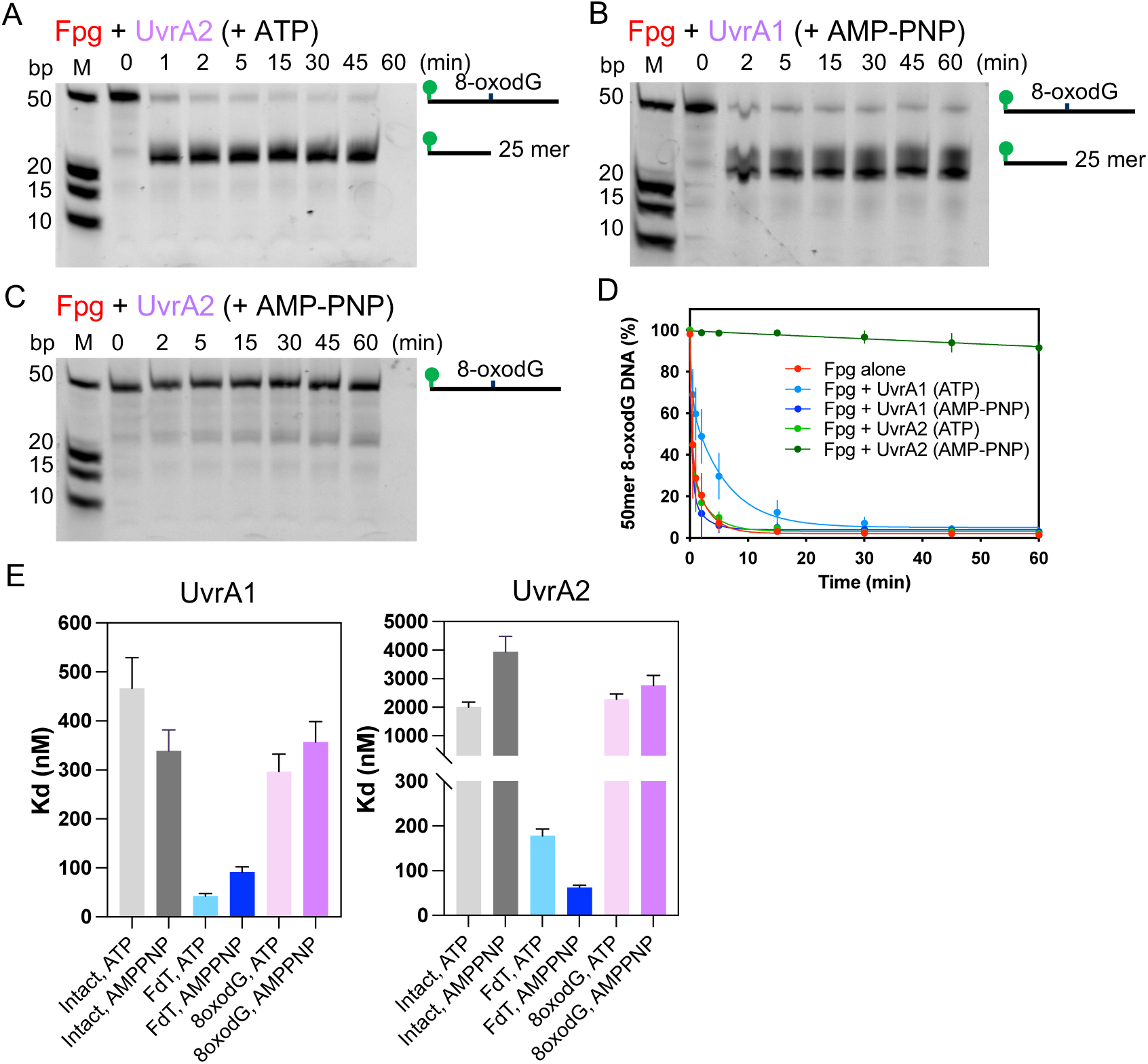
UvrA1 and UvrA2 cause delay in Fpg activity in a nucleotide-dependent manner. (A)-(C) Representative urea polyacrylamide gels illustrating the incision activity of 25 nM Fpg in the presence of 25 nM UvrA2 and ATP (A), 25 nM Fpg with 25 nM UvrA1 and AMP-PNPN (B) and 25 nM Fpg and 25 nM UvrA2 and AMP-PNP (C). (D) Kinetics of incision of 8-oxoG DNA by either Fpg alone (red), Fpg + UvrA1 in the presence of either ATP (light blue) or AMP-PNP (dark blue), and Fpg + UvrA2 in the presence of either ATP (light green) or AMP-PNP (dark green). Data points corresponding to the mean of three independent measurements were fitted to a two-phase exponential decay model. Error bars correspond to the standard deviation. (E) Binding constants (Kd) for UvrA1 (left) and UvrA2 (right) binding to intact 50mer dsDNA(grey), FdT-containing dsDNA (blue) and 8-oxoG-containing dsDNA (pink) in either the presence of ATP (light colors) or AMP-PNP (dark colors). The histograms represent the K_d_ values derived from at least three independent measurements and the error bars correspond to the standard error of the mean.

To determine whether specific binding of the UvrAs to 8-oxoG DNA in a nucleotide-dependent manner may impede Fpg activity, we performed fluorescence polarization experiments to assess the binding affinity of UvrA1 and UvrA2 to 8-oxoG containing DNA in the presence of either ATP or AMP-PNP (Fig. 6E and Fig. S7). For comparison, we also performed these measurements on intact DNA and a classical NER substrate, a DNA bearing a fluorescein-conjugated thymine (FdT) instead of the 8-oxoG nucleotide. Our results confirmed that indeed UvrA1 specifically binds to FdT DNA with enhanced affinity in the presence of ATP, while conversely, UvrA2 binds FdT DNA tightly in the presence of AMP-PNP (Fig. 6E). Both UvrA1 and UvrA2 also showed binding to the oxidized DNA substrate, but with K_d_ values 5-40-fold higher than for their classical NER substrate, and similar to that obtained for undamaged DNA (Fig. 6E and Fig. S7). The binding to undamaged and 8-oxoG DNA was only mildly affected by the nucleotide-bound state of the protein. A notable difference between UvrA1 and UvrA2 is their affinity for undamaged DNA. UvrA1 binds to undamaged and 8-oxoG DNA with an affinity in the order of 300-500 nM, while UvrA2 binds to these substrates with K_d_ values above 2 µM (Fig. 6E and Fig. S7). These results thus suggest that the UvrA proteins do not specifically recognize 8-oxoG in DNA, but rather bind non-specifically to this substrate in the presence of both ATP and AMP-PNP, which may contribute to the delayed Fpg activity.

## DISCUSSION

Traditional paradigms have long held that non-bulky DNA lesions resulting from oxidative stress are primarily repaired through the BER pathway, whereas bulky lesions are instead processed by the NER pathway. However, recent studies suggest that this dichotomy is overly simplistic, and that these pathways may cooperate or compete for the repair of specific types of DNA damage.

The present study provides the first experimental evidence for a complex functional interplay between the BER and NER pathways in *D. radiodurans*, occurring at three distinct levels (Fig. 7): (1) direct protein-protein interactions, (2) competitive modulation of repair activity, and (3) synergistic processing of repair intermediates. First, we identified direct physical interactions between components of the two pathways, notably involving both UvrA variants. We found that UvrA2 interacts with five DNA glycosylases—Fpg, EndoIII1, EndoIII2, EndoIII3, and UNG—responsible for the removal of oxidized bases and mis-incorporated uracil. In addition, UvrA1 was found to interact with Fpg and the DNA ligase A. Biophysical measurements further indicated that these interactions are weak and transient, with affinity constants in the low micromolar range. These findings identify the UvrA proteins—and particularly the UvrA2 variant—as key mediators of crosstalk between the NER and BER pathways. Unlike in *E. coli*, where a functional interplay between NER and BER has previously been reported (24, 25), here we provide evidence of direct physical interactions between these pathways, with UvrA2 acting as a central hub. Interestingly, UvrA2 was also recently detected in the interactomes of three DSB repair factors in *D. radiodurans* (DdrB, RecA, and SSB) (36), supporting a broader coordinating role for this protein in the DNA damage response in this radiation resistant organism.

**Figure 7.**
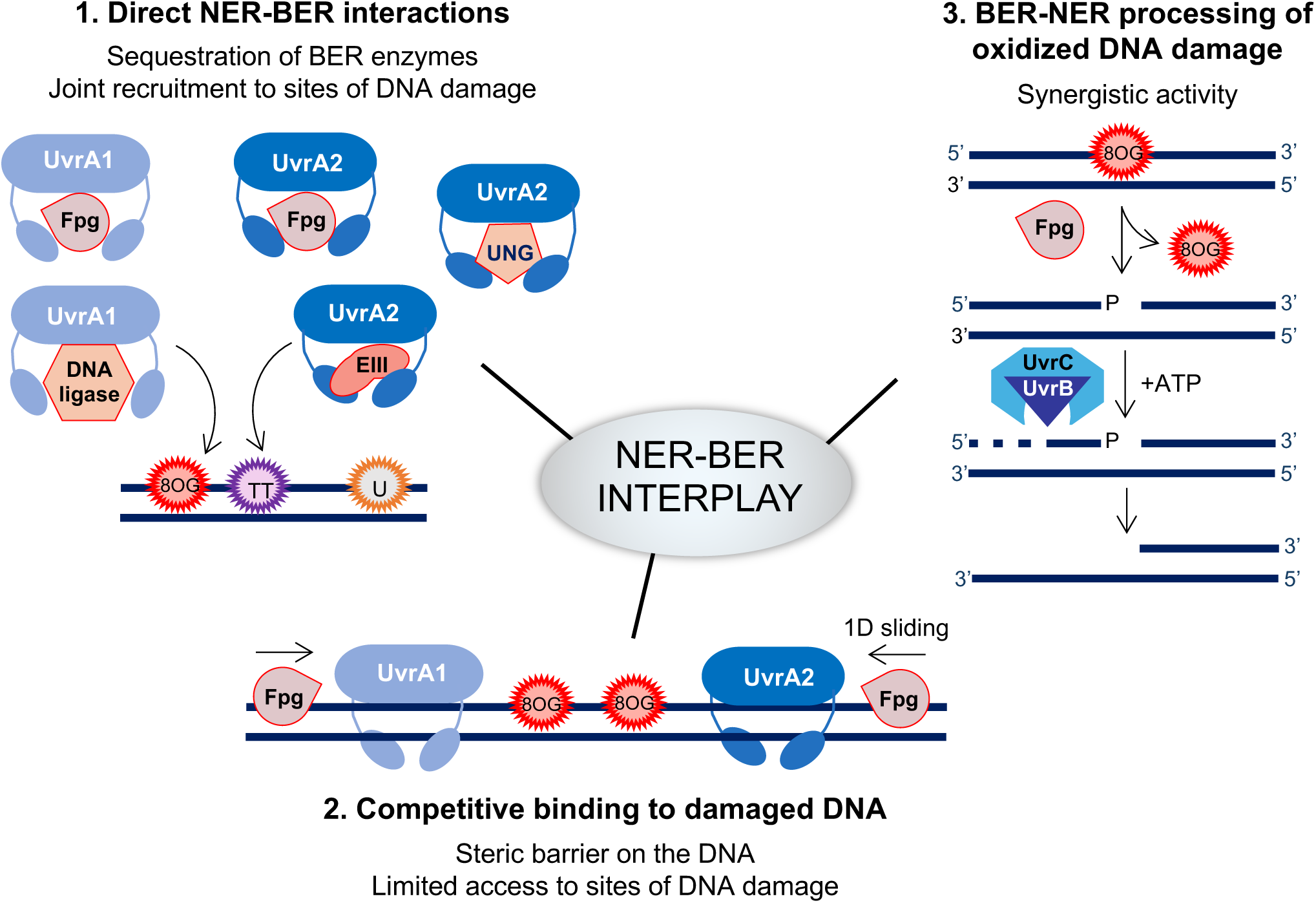
Interplay between BER and NER in *D. radiodurans*. Schematic diagram illustrating the three levels of interplay identified between BER and NER: (1) direct protein-protein interactions between UvrA1 or UvrA2 and BER enzymes, (2) competitive modulation of BER repair activity by NER factors, and (3) synergistic processing of repair intermediates by BER and NER pathways.

Direct NER-BER interactions may be a means of regulating their repair activity or instead promoting the co-recruitment of BER and NER machineries to favor coordinated repair of clustered DNA lesions (55). The delayed activity of Fpg in the presence of UvrA, and in particular its almost complete inhibition by UvrA2 in AMP-PNP conditions, together with the predicted AlphaFold3 models of UvrA–BER complexes support a model in which direct UvrA-BER interactions may maintain both systems in an inactive or resting state. In the predicted AlphaFold3 models of UvrA–BER complexes, the BER enzymes were indeed consistently predicted to bind in between the flexible IDs of UvrA—which function as “arms” to trap and probe the DNA duplex during damage recognition (42, 56–58). If confirmed experimentally, such an arrangement could sterically hinder both DNA binding by the UvrA variants and substrate engagement by the BER enzymes. Each UvrA monomer possesses two ATP binding sites that tightly regulate its dimerization, DNA scanning and damage recognition functions (42, 59, 60). It seems quite plausible that the nucleotide-bound state may also regulate UvrA’s binding to its cellular partners, notably by modulating the flexibility of the IDs, which are inserted within the N-terminal NBDs of UvrA factors (42, 57).

Beyond physical interactions, a second level of NER-BER interplay revealed in this work is the competition between NER and BER factors for the repair of oxidized bases, suggesting a potential regulatory mechanism for pathway selection (Fig. 7). The Uvr proteins were indeed found to slow down to varying extents the repair of 8-oxoG by Fpg. A similar delay in the repair of hydantoins by NEIL1 was observed *in vitro* in the presence of the eukaryotic NER damage sensor complex XPC-RAD23B (61), which was proposed to result from competitive binding of XPC-RAD23B to the site of DNA damage, thereby preventing incision by NEIL1, and not from altered NEIL1 reaction kinetics. A competitive relationship between NER and BER factors has also been reported previously in human cell extracts for the repair of DNA hydantoin lesions resulting from oxidation of 8-oxoG (20–23). In these studies, Shafirovich and colleagues showed that such lesions can be processed either by the NER or the BER pathways, producing two distinct molecular signatures (dual versus single incision). Importantly, as in the XPC-RAD23B study, we do not observe any processing of the 8-oxoG DNA by the NER system alone.

Interestingly, in the case of the UvrA-Fpg interplay in *D. radiodurans*, our fluorescence polarization data strongly indicates that specific binding of UvrA1 or UvrA2 to the 8-oxoG sites is not the main cause of delayed Fpg activity. Instead, the observed inhibition of Fpg activity may be due to direct interactions between Fpg and UvrAs, as discussed above, but also to non-specific DNA binding by the Uvr factors that physically obstruct Fpg’s displacement along the DNA, rather than competitive binding to the oxidized base. Consistent with this, we found that the full NER machinery (UvrA1–UvrB–UvrC), and to a lesser extent the UvrB/UvrC complex alone, significantly delayed Fpg-mediated repair of oxidized DNA. In the absence of ATP, when UvrB/UvrC stably associates with DNA, inhibition was even stronger, leaving a larger fraction of uncleaved DNA. These findings support a model in which NER components can transiently block access of BER enzymes to damaged sites through non-specific DNA occupancy.

Finally, a third level of NER-BER crosstalk was identified in this study, involving a synergistic processing of DNA damage by BER and NER factors (Fig. 7). We notably show that the UvrB/UvrC factors can further process the product of the Fpg incision (β- and δ-elimination) reaction, *i.e.* 1-nt gapped dsDNA with a single-strand break in place of the 8-oxoG base. This activity is independent of the DNA damage sensor, UvrA1, but is, nonetheless, ATP-driven and carried out by the RNase H domain of UvrC, resulting in the accumulation of multiple short DNA fragments, notably 7 mer fragments, arising from specific cleavage of the DNA fragment located on the 5’ side of the gap. It should be noted that we have previously shown that UvrB/UvrC does not incise the classical NER substrate (FdT containing DNA) or even 3′ incised DNA in this way (44), indicating that this activity is specific to certain DNA structures or termini. Although this is, to the best of our knowledge, the first report of bacterial NER directly processing the gapped DNA produced by a BER bifunctional DNA glycosylase, there have been earlier reports, notably on *E. coli* UvrB/UvrC, demonstrating that UvrB/UvrC can act as a structure-specific nuclease, capable of incising DNA bearing pre-existing nicks, gaps or 5’ overhangs a few nucleotides 5′ of the nick or gap thereby releasing reproducible cleavage products (62, 63). Similarly, this activity was shown to be UvrA-independent and ATP-dependent, relying on the ATP-driven helicase activity of UvrB, which is stimulated by both UvrC binding and DNA binding. Beyond its canonical NER activity on UV-induced or bulky DNA lesions, *D. radiodurans*, like *E. coli*, thus exhibits an additional UvrB/UvrC-dependent repair activity that targets specific DNA structures and possibly distortions, rather than DNA damage itself. This dual functionality of UvrB/UvrC may be particularly critical in *D. radiodurans*, where high doses of ionizing radiation generate clustered lesions (55). By rapidly processing BER intermediates, *D. radiodurans* could minimize the accumulation of toxic single-strand breaks, thereby enhancing genomic stability under extreme stress. Of course, how frequently bacterial cells will route a BER intermediate into an NER-like processing pathway *in vivo* will most likely depend on timing, relative protein abundance, and whether the intermediate is exposed long enough. These findings also suggest that beyond the processing of BER repair intermediates, the NER factors UvrB/UvrC may be able to process intermediates produced by other repair pathways, thereby modulating the choice of the downstream repair pathway.

Together, these findings support a model in which *D. radiodurans* integrates multiple repair systems to preserve genome integrity under extreme stress. Future *in vivo* studies will be needed to clarify the functional relevance of these mechanisms and determine if and how they may contribute to this organism’s exceptional resilience to DNA damage.

## ACKNOWLEDGEMENTS

IBS acknowledges integration into the Interdisciplinary Research Institute of Grenoble (IRIG, CEA). This work used the biophysics platform of the Grenoble Instruct-ERIC center (ISBG; UMS 3518 CNRS-CEA-UGA-EMBL) within the Grenoble Partnership for Structural Biology (PSB), supported by FRISBI (ANR-10-INBS-0005-02) and the GRAL and ARCANE labex, financed within the University Grenoble Alpes graduate school (Ecoles Universitaires de Recherche) CBH-EUR-GS (ANR-17-EURE-0003, ANR-15-IDEX-02).

## DATA AVAILABILITY

The data underlying this article are available in the article and in its online supplementary material.

## REFERENCES

1. Chatterjee, N. and Walker, G.C. (2017) Mechanisms of DNA damage, repair and mutagenesis. Environ Mol Mutagen, 58, 235–263.

2. Barnes, J.L., Zubair, M., John, K., Poirier, M.C. and Martin, F.L. (2018) Carcinogens and DNA damage. Biochemical Society Transactions, 46, 1213–1224.

3. Lindahl, T. and Barnes, D.E. (2000) Repair of endogenous DNA damage. Cold Spring Harb Symp Quant Biol, 65, 127–133.

4. Tubbs, A. and Nussenzweig, A. (2017) Endogenous DNA Damage as a Source of Genomic Instability in Cancer. Cell, 168, 644–656.

5. Lindahl, T. (1993) Instability and decay of the primary structure of DNA. Nature, 362, 709–715.

6. Torgovnick, A. and Schumacher, B. (2015) DNA repair mechanisms in cancer development and therapy. Front Genet, 6, 157.

7. Huang, R. and Zhou, P.-K. (2021) DNA damage repair: historical perspectives, mechanistic pathways and clinical translation for targeted cancer therapy. Signal Transduct Target Ther, 6, 254.

8. Maréchal, A. and Zou, L. (2013) DNA Damage Sensing by the ATM and ATR Kinases. Cold Spring Harb Perspect Biol, 5, a012716.

9. Wozniak, K.J. and Simmons, L.A. (2022) Bacterial DNA excision repair pathways. Nat Rev Microbiol, 20, 465–477.

10. Oh, J.-M., Kang, Y., Park, J., Sung, Y., Kim, D., Seo, Y., Lee, E.A., Ra, J.S., Amarsanaa, E., Park, Y.-U., et al. (2023) MSH2-MSH3 promotes DNA end resection during homologous recombination and blocks polymerase theta-mediated end-joining through interaction with SMARCAD1 and EXO1. Nucleic Acids Res, 51, 5584–5602.

11. Oh, J.-M. and Myung, K. (2022) Crosstalk between different DNA repair pathways for DNA double strand break repairs. Mutation Research/Genetic Toxicology and Environmental Mutagenesis, 873, 503438.

12. Kumar, N., Theil, A.F., Roginskaya, V., Ali, Y., Calderon, M., Watkins, S.C., Barnes, R.P., Opresko, P.L., Pines, A., Lans, H., et al. (2022) Global and transcription-coupled repair of 8-oxoG is initiated by nucleotide excision repair proteins. Nat Commun, 13, 974.

13. Kumar, N., Raja, S. and Van Houten, B. (2020) The involvement of nucleotide excision repair proteins in the removal of oxidative DNA damage. Nucleic Acids Research, 48, 11227–11243.

14. Jang, S., Kumar, N., Beckwitt, E.C., Kong, M., Fouquerel, E., Rapić-Otrin, V., Prasad, R., Watkins, S.C., Khuu, C., Majumdar, C., et al. (2019) Damage sensor role of UV-DDB during base excision repair. Nature structural & molecular biology, 26, 695–703.

15. Jang, S., Schaich, M.A., Khuu, C., Schnable, B.L., Majumdar, C., Watkins, S.C., David, S.S. and Van Houten, B. (2021) Single molecule analysis indicates stimulation of MUTYH by UV-DDB through enzyme turnover. Nucleic Acids Research, 49, 8177–8188.

16. D’Errico, M., Parlanti, E., Teson, M., de Jesus, B.M.B., Degan, P., Calcagnile, A., Jaruga, P., Bjørås, M., Crescenzi, M., Pedrini, A.M., et al. (2006) New functions of XPC in the protection of human skin cells from oxidative damage. EMBO J, 25, 4305–4315.

17. Menoni, H., Wienholz, F., Theil, A.F., Janssens, R.C., Lans, H., Campalans, A., Radicella, J.P., Marteijn, J.A. and Vermeulen, W. (2018) The transcription-coupled DNA repair-initiating protein CSB promotes XRCC1 recruitment to oxidative DNA damage. Nucleic Acids Res, 46, 7747–7756.

18. Melis, J.P.M., van Steeg, H. and Luijten, M. (2013) Oxidative DNA damage and nucleotide excision repair. Antioxid Redox Signal, 18, 2409–2419.

19. Parlanti, E., D’Errico, M., Degan, P., Calcagnile, A., Zijno, A., van der Pluijm, I., van der Horst, G.T.J., Biard, D.S.F. and Dogliotti, E. (2012) The cross talk between pathways in the repair of 8-oxo-7, 8-dihydroguanine in mouse and human cells. Free Radic Biol Med, 53, 2171–2177.

20. Shafirovich, V. and Geacintov, N.E. (2017) Removal of oxidatively generated DNA damage by overlapping repair pathways. Free Radic Biol Med, 107, 53–61.

21. Shafirovich, V. and Geacintov, N.E. (2021) Excision of Oxidatively Generated Guanine Lesions by Competitive DNA Repair Pathways. Int J Mol Sci, 22.

22. Shafirovich, V., Kropachev, K., Kolbanovskiy, M. and Geacintov, N.E. (2019) Excision of Oxidatively Generated Guanine Lesions by Competing Base and Nucleotide Excision Repair Mechanisms in Human Cells. Chem Res Toxicol, 32, 753–761.

23. Shafirovich, V., Kropachev, K., Anderson, T., Liu, Z., Kolbanovskiy, M., Martin, B.D., Sugden, K., Shim, Y., Chen, X., Min, J.-H., et al. (2016) Base and Nucleotide Excision Repair of Oxidatively Generated Guanine Lesions in DNA *. Journal of Biological Chemistry, 291, 5309–5319.

24. Czeczot, H., Tudek, B., Lambert, B., Laval, J. and Boiteux, S. (1991) *Escherichia coli* Fpg protein and UvrABC endonuclease repair DNA damage induced by methylene blue plus visible light in vivo and in vitro. J Bacteriol, 173, 3419–3424.

25. Asad, N.R., De Almeida, C.E.B., Asad, L.M.B.O., Felzenszwalb, I. and Leitão, A.C. (1995) Fpg and UvrA proteins participate in the repair of DNA lesions induced by hydrogen peroxide in low iron level in *Escherichia coli*. Biochimie, 77, 262–264.

26. Couvé-Privat, S., Macé, G., Rosselli, F. and Saparbaev, M.K. (2007) Psoralen-induced DNA adducts are substrates for the base excision repair pathway in human cells. Nucleic Acids Res, 35, 5672–5682.

27. Couvé, S., Macé-Aimé, G., Rosselli, F. and Saparbaev, M.K. (2009) The human oxidative DNA glycosylase NEIL1 excises psoralen-induced interstrand DNA cross-links in a three-stranded DNA structure. J Biol Chem, 284, 11963–11970.

28. Pr, M., S, C., C, Z., Ms, A., R, G., Bt, M., Jl, P., Rh, E. and Mk, S. (2017) The Human DNA glycosylases NEIL1 and NEIL3 Excise Psoralen-Induced DNA-DNA Cross-Links in a Four-Stranded DNA Structure. Scientific reports, 7.

29. Mullins, E.A., Rodriguez, A.A., Bradley, N.P. and Eichman, B.F. (2019) Emerging Roles of DNA Glycosylases and the Base Excision Repair Pathway. Trends Biochem Sci, 44, 765–781.

30. Wilson, D.M. and Seidman, M.M. (2010) A novel link to base excision repair? Trends Biochem Sci, 35, 247–252.

31. Lim, S., Jung, J.-H., Blanchard, L. and de Groot, A. (2018) Conservation and diversity of radiation and oxidative stress resistance mechanisms in *Deinococcus* species. FEMS Microbiol Rev, 43, 19–52.

32. Liu, F., Li, N. and Zhang, Y. (2023) The radioresistant and survival mechanisms of *Deinococcus radiodurans*. Radiation Medicine and Protection, 4, 70–79.

33. Slade, D. and Radman, M. (2011) Oxidative Stress Resistance in *Deinococcus radiodurans*. Microbiol Mol Biol Rev, 75, 133–191.

34. Cox, M.M. and Battista, J.R. (2005) *Deinococcus radiodurans* - the consummate survivor. Nat Rev Microbiol, 3, 882–892.

35. Timmins, J. and Moe, E. (2016) A Decade of Biochemical and Structural Studies of the DNA Repair Machinery of *Deinococcus radiodurans:* Major Findings, Functional and Mechanistic Insight and Challenges. Computational and Structural Biotechnology Journal, 14, 168–176.

36. Ujaoney, A.K., Padwal, M.K. and Basu, B. (2021) An *in vivo* Interaction Network of DNA-Repair Proteins: A Snapshot at Double Strand Break Repair in *Deinococcus radiodurans*. J. Proteome Res., 20, 3242–3255.

37. Krisko, A. and Radman, M. (2013) Biology of Extreme Radiation Resistance: The Way of *Deinococcus radiodurans*. Cold Spring Harb Perspect Biol, 5, a012765.

38. Tanaka, M., Narumi, I., Funayama, T., Kikuchi, M., Watanabe, H., Matsunaga, T., Nikaido, O. and Yamamoto, K. (2005) Characterization of pathways dependent on the *uvsE*, *uvrA1*, or *uvrA2* gene product for UV resistance in *Deinococcus radiodurans*. J Bacteriol, 187, 3693–3697.

39. Zahradka, K., Slade, D., Bailone, A., Sommer, S., Averbeck, D., Petranovic, M., Lindner, A.B. and Radman, M. (2006) Reassembly of shattered chromosomes in *Deinococcus radiodurans*. Nature, 443, 569–573.

40. Seck, A., De Bonis, S., Saint-Pierre, C., Gasparutto, D., Ravanat, J.-L. and Timmins, J. (2022) In vitro reconstitution of an efficient nucleotide excision repair system using mesophilic enzymes from *Deinococcus radiodurans*. Commun Biol, 5, 127.

41. Makarova, K.S., Aravind, L., Wolf, Y.I., Tatusov, R.L., Minton, K.W., Koonin, E.V. and Daly, M.J. (2001) Genome of the Extremely Radiation-Resistant Bacterium *Deinococcus radiodurans* Viewed from the Perspective of Comparative Genomics. Microbiology and Molecular Biology Reviews, 65, 44–79.

42. Timmins, J., Gordon, E., Caria, S., Leonard, G., Acajjaoui, S., Kuo, M.S., Monchois, V. and McSweeney, S. (2009) Structural and mutational analyses of *Deinococcus radiodurans* UvrA2 provide insight into DNA binding and damage recognition by UvrAs. Structure, 17, 547–58.

43. Battesti, A. and Bouveret, E. (2012) The bacterial two-hybrid system based on adenylate cyclase reconstitution in *Escherichia coli*. Methods, 58, 325–34.

44. Seck, A., De Bonis, S., Stelter, M., Ökvist, M., Senarisoy, M., Hayek, M.R., Le Roy, A., Martin, L., Saint-Pierre, C., Silveira, C.M., et al. (2023) Structural and functional insights into the activation of the dual incision activity of UvrC, a key player in bacterial NER. Nucleic Acids Res, 51, 2931–2949.

45. Sarre, A., Ökvist, M., Klar, T., Hall, D.R., Smalås, A.O., McSweeney, S., Timmins, J. and Moe, E. (2015) Structural and functional characterization of two unusual endonuclease III enzymes from *Deinococcus radiodurans*. J Struct Biol, 191, 87–99.

46. Sarre, A., Stelter, M., Rollo, F., De Bonis, S., Seck, A., Hognon, C., Ravanat, J.-L., Monari, A., Dehez, F., Moe, E., et al. (2019) The three Endonuclease III variants of *Deinococcus radiodurans* possess distinct and complementary DNA repair activities. DNA Repair (Amst), 78, 45–59.

47. Leiros, I., Moe, E., Smalås, A.O. and McSweeney, S. (2005) Structure of the uracil-DNA N-glycosylase (UNG) from *Deinococcus radiodurans*. Acta Crystallogr D Biol Crystallogr, 61, 1049–1056.

48. Fernandes, A., Williamson, A., Matias, P.M. and Moe, E. (2023) Structure/function studies of the NAD+-dependent DNA ligase from the poly-extremophile *Deinococcus radiodurans* reveal importance of the BRCT domain for DNA binding. Extremophiles, 27, 26.

49. Evans, R., O’Neill, M., Pritzel, A., Antropova, N., Senior, A., Green, T., Žídek, A., Bates, R., Blackwell, S., Yim, J., et al. (2021) Protein complex prediction with AlphaFold-Multimer. 10.1101/2021.10.04.463034.

50. Abramson, J., Adler, J., Dunger, J., Evans, R., Green, T., Pritzel, A., Ronneberger, O., Willmore, L., Ballard, A.J., Bambrick, J., et al. (2024) Accurate structure prediction of biomolecular interactions with AlphaFold 3. Nature, 630, 493–500.

51. Krissinel, E. and Henrick, K. (2007) Inference of macromolecular assemblies from crystalline state. J Mol Biol, 372, 774–797.

52. Pettersen, E.F., Goddard, T.D., Huang, C.C., Meng, E.C., Couch, G.S., Croll, T.I., Morris, J.H. and Ferrin, T.E. (2021) UCSF ChimeraX: Structure visualization for researchers, educators, and developers. Protein Science, 30, 70–82.

53. Kasai, H. and Nishimura, S. (1984) Hydroxylation of deoxyguanosine at the C-8 position by ascorbic acid and other reducing agents. Nucleic Acids Research, 12, 2137–2145.

54. Koval, V.V. (2004) Pre-steady-state kinetics shows differences in processing of various DNA lesions by Escherichia coli formamidopyrimidine-DNA glycosylase. Nucleic Acids Research, 32, 926–935.

55. Sage, E. and Shikazono, N. (2017) Radiation-induced clustered DNA lesions: Repair and mutagenesis. Free Radical Biology and Medicine, 107, 125–135.

56. Gantchev, T.G. and Hunting, D.J. (2010) Modeling the interactions of the nucleotide excision repair UvrA(2) dimer with DNA. Biochemistry, 49, 10912–24.

57. Pakotiprapha, D., Inuzuka, Y., Bowman, B.R., Moolenaar, G.F., Goosen, N., Jeruzalmi, D. and Verdine, G.L. (2008) Crystal structure of *Bacillus stearothermophilus* UvrA provides insight into ATP-modulated dimerization, UvrB interaction, and DNA binding. Mol Cell, 29, 122–33.

58. Jaciuk, M., Nowak, E., Skowronek, K., Tanska, A. and Nowotny, M. (2011) Structure of UvrA nucleotide excision repair protein in complex with modified DNA. Nat Struct Mol Biol, 18, 191–7.

59. Barnett, J.T. and Kad, N.M. (2019) Understanding the coupling between DNA damage detection and UvrA’s ATPase using bulk and single molecule kinetics. The FASEB Journal, 33, 763–769.

60. Case, B.C., Hartley, S., Osuga, M., Jeruzalmi, D. and Hingorani, M.M. (2019) The ATPase mechanism of UvrA2 reveals the distinct roles of proximal and distal ATPase sites in nucleotide excision repair. Nucleic Acids Res, 47, 4136–4152.

61. Kolbanovskiy, M., Shim, Y., Min, J.-H., Geacintov, N.E. and Shafirovich, V. (2020) Inhibition of Excision of Oxidatively Generated Hydantoin DNA Lesions by NEIL1 by the Competitive Binding of the Nucleotide Excision Repair Factor XPC-RAD23B. Biochemistry, 59, 1728–1736.

62. Moolenaar, G.F., Bazuine, M., van Knippenberg, I.C., Visse, R. and Goosen, N. (1998) Characterization of the *Escherichia coli* damage-independent UvrBC endonuclease activity. J Biol Chem, 273, 34896–34903.

63. Zou, Y., Walker, R., Bassett, H., Geacintov, N.E. and Van Houten, B. (1997) Formation of DNA repair intermediates and incision by the ATP-dependent UvrB-UvrC endonuclease. J Biol Chem, 272, 4820–7.

