## Supplemental Figures and Tables for "Evidence for strong interplay between the nucleotide and base excision repair pathways in *D. radiodurans*"

Supplementary Figures S1-S7
Supplementary Tables S1-S2

### Figure S1

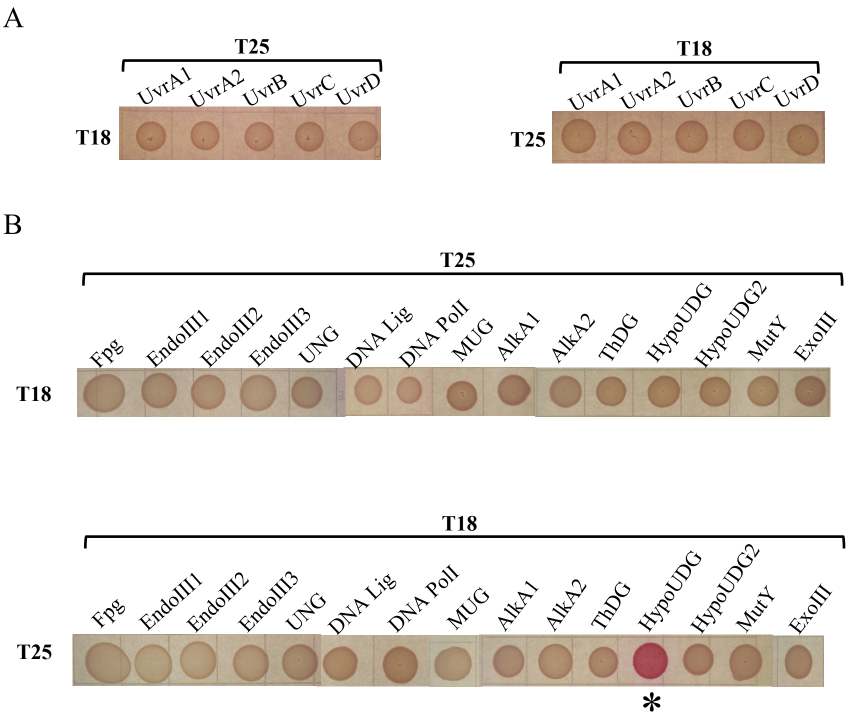


**Figure S1. Single negative controls of all NER & BER constructs.** (A) Single negative control of all NER proteins in both combinations. (B) Single negative control of all BER proteins in both combinations.

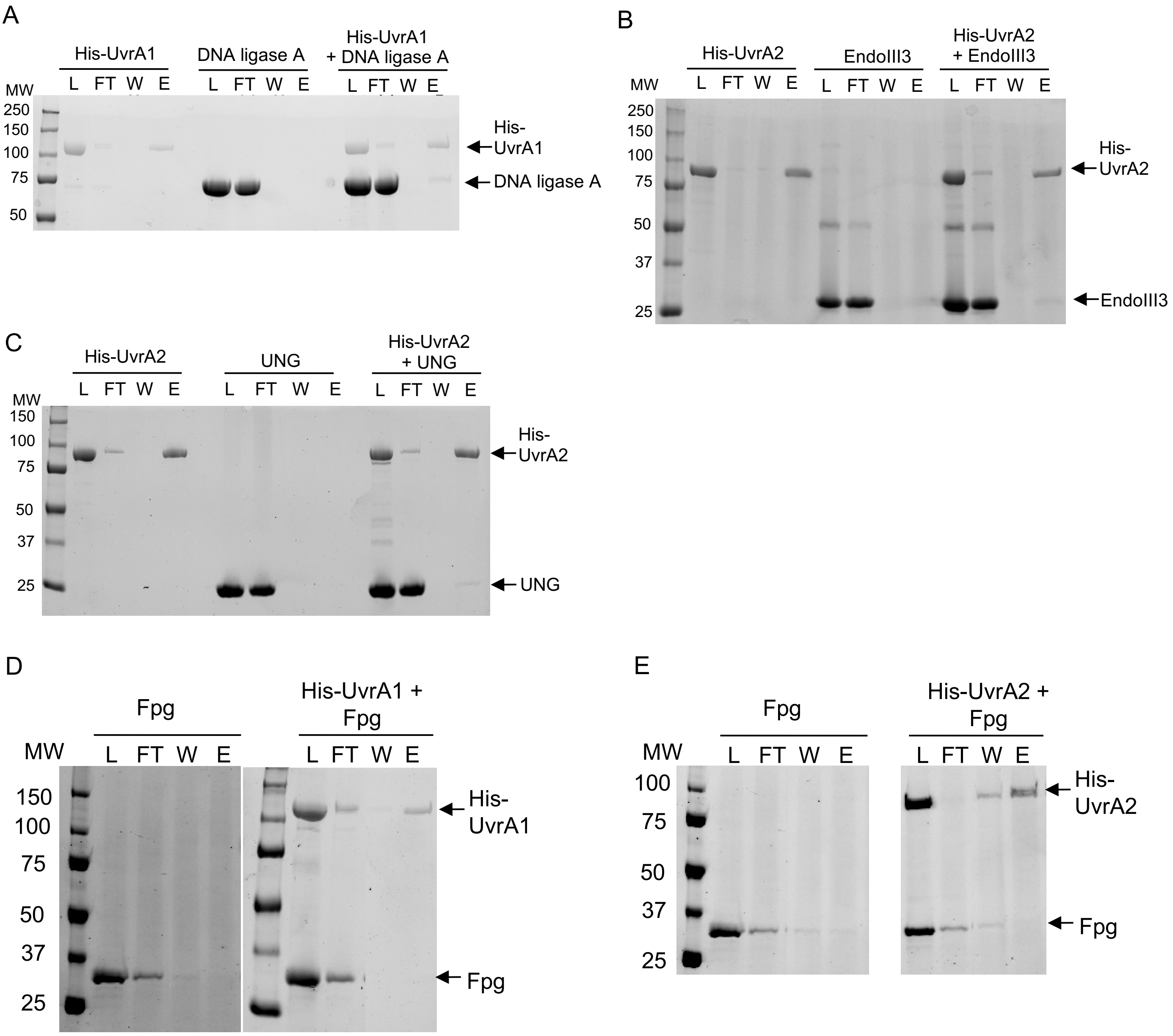


**Figure S2: Pull-down analysis of UvrA-DNA glycosylase interactions.** (A)-(E) SDS-PAGE analysis of the pull-down experiments performed to evaluate the binding of His-tagged UvrA1 (A and D) or His-tagged UvrA2 (B, C and E) to DNA ligase A (A), EndoIII3 (B), UNG (C) or Fpg (D-E). (A-C) In each case, the first two experiments (left and middle) are control experiments to verify the specificity of the binding to the Nickel affinity resin and to the His-tagged bait. Right: His-tagged UvrA1 (A, D) or UvrA2 (B, C and E) bait was mixed with a 5-fold molar excess of non-tagged prey protein and loaded (L) on Nickel affinity resin. The flow-through (FT) and the final wash (W) were collected before the protein was eluted (E). Molecular weight (MW) markers are indicated in kDa.

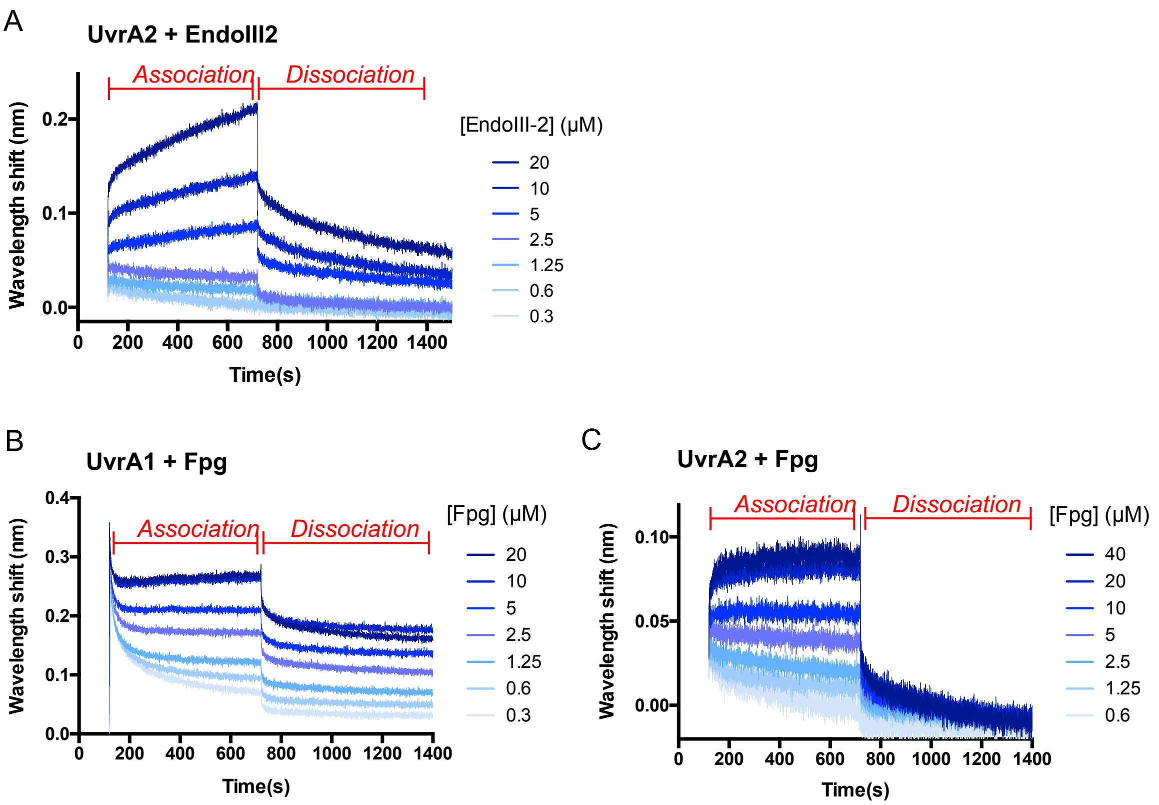


**Figure S3: BLI measurements of UvrA-DNA glycosylase interactions.** (A) Real-time sensorgram traces reflecting changes in wavelength shift during the association and dissociation of 0.3 to 20 μM EndoIII2 (light to dark blue) on immobilized biotinylated UvrA2. (B) Real-time sensorgram traces reflecting changes in wavelength shift during the association and dissociation of 0.3 to 20 μM Fpg (light to dark blue) on immobilized biotinylated UvrA1. (C) Real-time sensorgram traces reflecting changes in wavelength shift during the association and dissociation of 0.6 to 40 μM Fpg (light to dark blue) on immobilized biotinylated UvrA2.

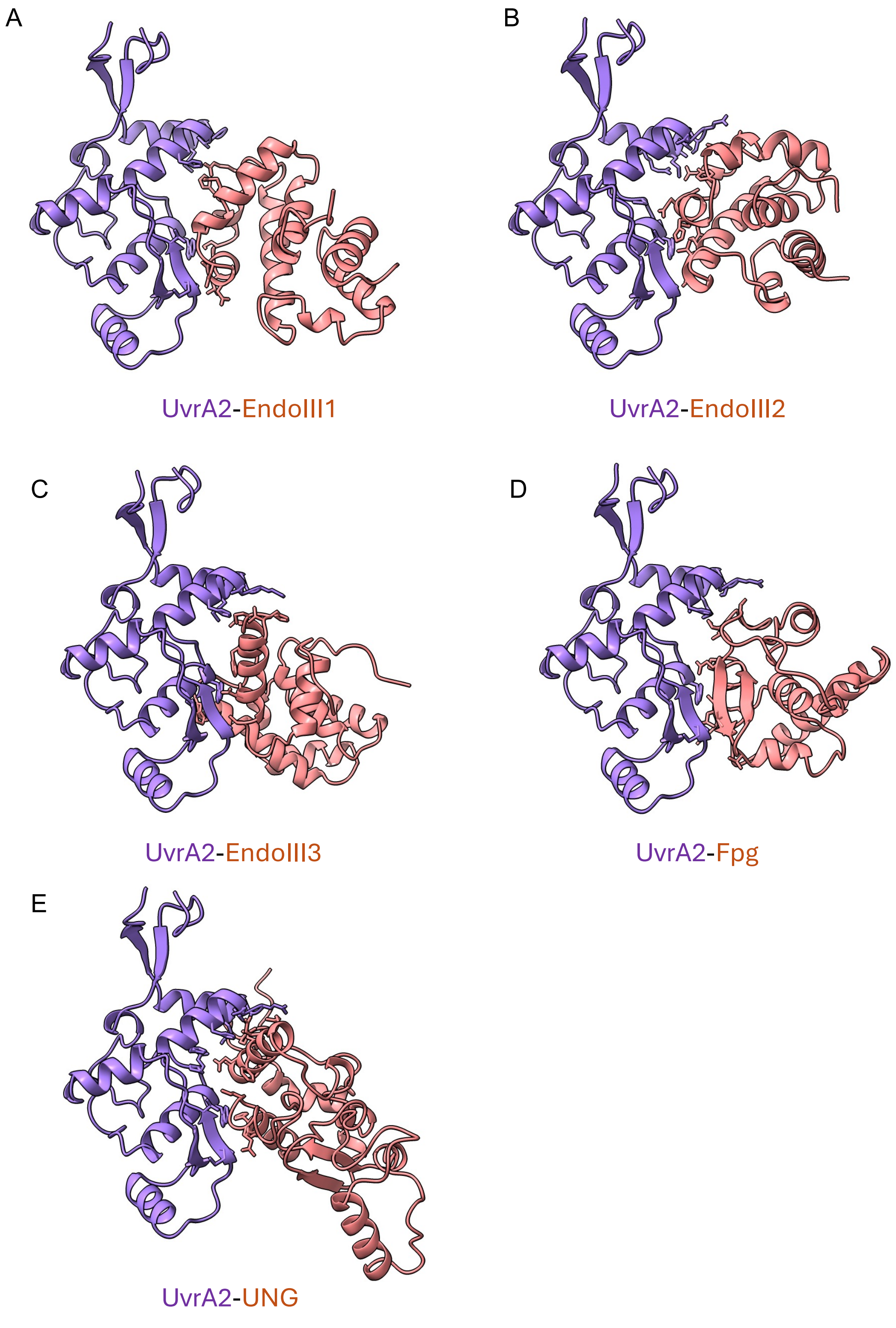


**Figure S4.** AlphaFold3 predicted models of UvrA2 insertion domains (purple) bound to BER factors (red): EndoIII1 (A), EndoIII2 (B), EndoIII3 (C), Fpg (D) and UNG (E). The illustrated models correspond to the top ranked AF3 complexes among the 5 models produced by AF3, with the highest ipTM scores (Table S2). Residues involved in the interaction interface are shown in sticks.

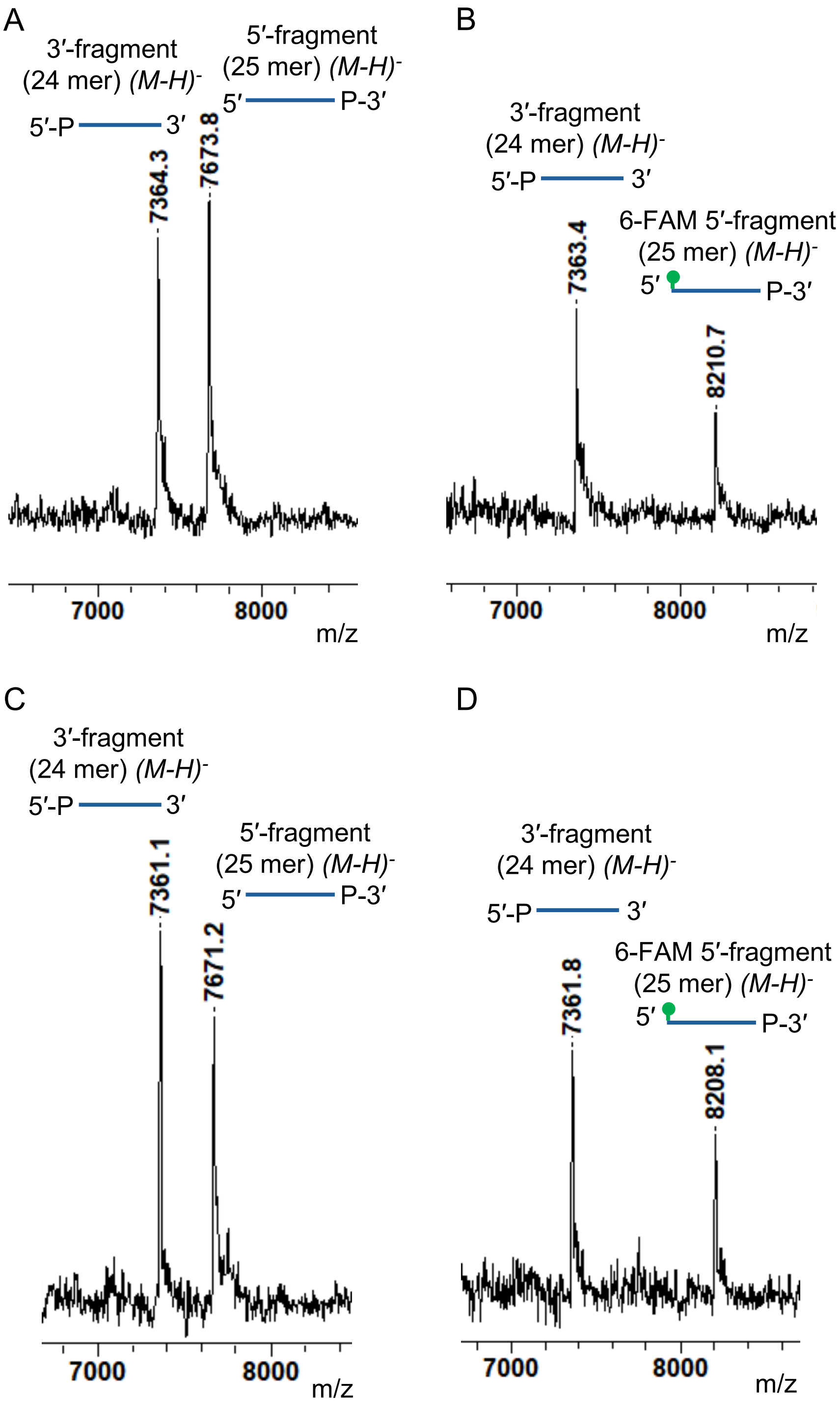


**Figure S5: MALDI-ToF analysis of Fpg treated DNA.** (A)-(B) MALDI-ToF MS spectra of the reaction products of 25 nM unlabeled (A) or 5'-FAM labeled (B) 50mer-8oxoG dsDNA treated with 25 nM Fpg for 60 min. Peaks corresponding to the 5' and 3' fragments released by Fpg processing of the 8-oxoG containing strand were observed in both cases. (C)-(D) MALDI-ToF MS spectra of 25 nM unlabeled (C) or 5'-FAM labeled (D) 50mer-gapped dsDNA. Peaks corresponding to the 5' and 3' fragments on either side of the 1-nt gap were detected at their expected molecular weights.

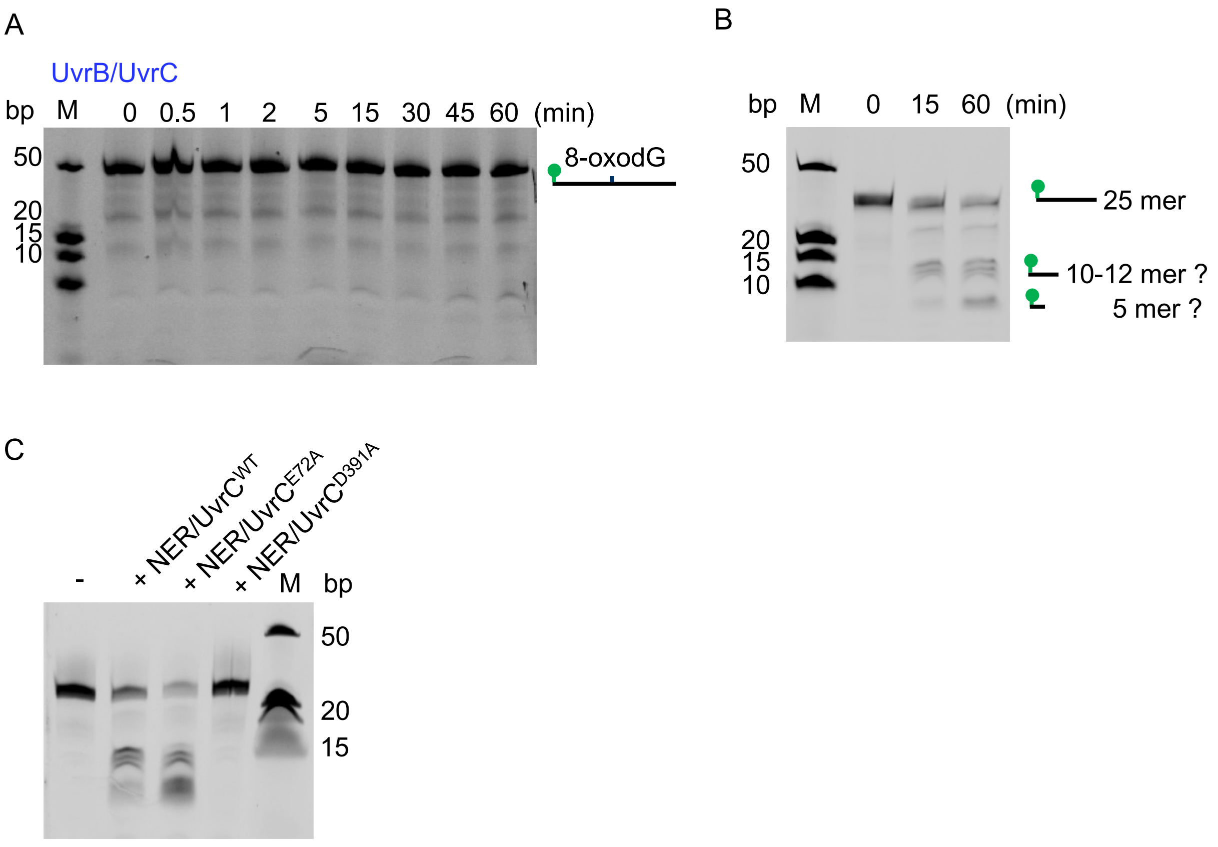


**Figure S6: Incision activity of the NER machinery on 8-oxoG or gapped dsDNA.** (A) Urea polyacrylamide gel showing the absence of incision of the 6-FAM labeled 50mer 8-oxoG substrate by the UvrB/UvrC proteins in the absence of Fpg. (B) Urea polyacrylamide gel showing the processing of 6-FAM labeled 50mer gapped (1-nt) dsDNA substrate by the UvrA1/UvrB/UvrC proteins after 0, 15 and 60 minutes of reaction. The fragments obtained are the same as those obtained when combining Fpg and the NER machinery of 8-oxoG dsDNA. (C) Urea polyacrylamide gel showing the processing of 6-FAM labeled 50mer gapped (1-nt) dsDNA substrate by UvrA1/UvrB supplemented with either wild-type (WT) $UvrC^{WT}$ or two point mutants ($UvrC^{E72A}$ or $UvrC^{D391A}$) after 60 minutes of reaction. Left lane: control experiment without any enzyme. The fragments obtained with the $UvrC^{E72A}$ mutant are the same as those obtained with $UvrC^{WT}$, whereas no cleavage was seen with $UvrC^{D391A}$.

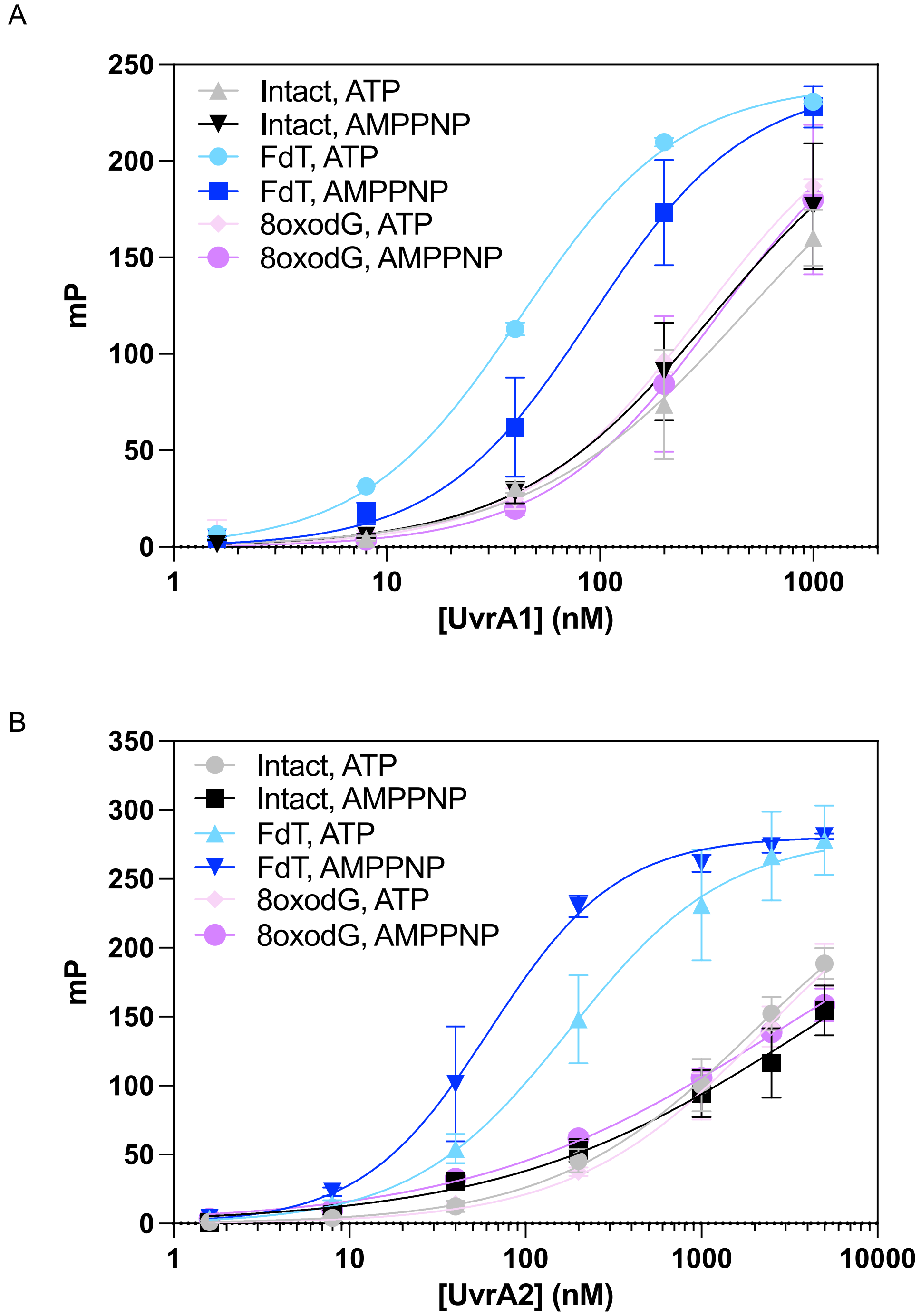


**Figure S7:** Fluorescence polarization titration curves of UvrA1 and UvrA2. 0 to 5 µM UvrA1 (A) or UvrA2 (B) were incubated with 2 nM 6-FAM labeled intact (grey/black) or lesion-containing dsDNA. FdT: fluorescein-conjugated thymine, a known NER substrate (light and dark blue). 8oxodG: 8-oxoguanine, a known BER/Fpg substrate (pink/purple). The reactions were performed in the presence of either ATP or the non-hydrolysable analogue, AMP-PNP. Data points are the mean of 4 independent measurements and the error bars correspond to the standard deviation. The data points were fitted to a one-site specific binding model with Hill slope in Graph Pad Prism 10.

### Table S1: Prediction scores and interface analysis of AlphaFold3 predicted models of UvrA-BER complexes

| UvrA variant | Partner | Ligand | Rank | Overall ipTM | Overall pTM | Ranking score | UvrA pTM (chains 1 & 2) | Partner pTM | ipTM (chains 1 & 2) | UvrA interacting region | Interacting area (chains 1 & 2) | Nhb/Nsb (chains 1 & 2) |
|---|---|---|---|---|---|---|---|---|---|---|---|---|
| UvrA1 | Fpg | 4xATP | 1 | 0.56 | 0.63 | 0.59 | 0.68/0.68 | 0.63 | 0.16/0.16 | UvrB-binding domain | 299/672 | 2; 2/4; 4 |
| UvrA1 | Fpg | 4xATP | 2 | 0.55 | 0.63 | 0.58 | 0.67/0.67 | 0.62 | 0.16/0.16 | NBDs | 614/739 | 6; 3/7; 6 |
| UvrA1 | Fpg | 4xATP | 3 | 0.54 | 0.62 | 0.57 | 0.68/0.67 | 0.63 | 0.21/0.16 | IDs | 530/1030 | 5; 3/8; 9 |
| UvrA1 | Fpg | 4xATP | 4 | 0.54 | 0.62 | 0.57 | 0.67/0.68 | 0.60 | 0.17/0.16 | IDs | 802/1273 | 4; 9/12; 7 |
| UvrA1 | Fpg | 4xATP | 5 | 0.53 | 0.61 | 0.56 | 0.67/0.67 | 0.63 | 0.17/0.17 | IDs | 121/638 | 0; 0/3; 6 |
| UvrA1 | DNA ligase | 4xATP | 1 | 0.49 | 0.58 | 0.52 | 0.67/0.67 | 0.52 | 0.17/0.17 | NBDs + UvrB-binding domain | 881/1337 | 5; 6/10; 8 |
| UvrA1 | DNA ligase | 4xATP | 2 | 0.48 | 0.58 | 0.52 | 0.69/0.68 | 0.52 | 0.17/0.17 | NBDs + IDs | 821/960 | 1; 6/6; 1 |
| UvrA1 | DNA ligase | 4xATP | 3 | 0.48 | 0.58 | 0.52 | 0.69/0.69 | 0.52 | 0.17/0.18 | IDs | 1175/1361 | 5; 10/7; 8 |
| UvrA1 | DNA ligase | 4xATP | 4 | 0.48 | 0.58 | 0.52 | 0.69/0.68 | 0.52 | 0.17/0.17 | NBDs + IDs | 687/833 | 2; 2/7; 4 |
| UvrA1 | DNA ligase | 4xATP | 5 | 0.47 | 0.57 | 0.51 | 0.68/0.68 | 0.53 | 0.17/0.18 | IDs | 935/1247 | 6; 3/10; 11 |
| UvrA2 | Fpg | 4xATP | 1 | 0.60 | 0.68 | 0.62 | 0.76/0.76 | 0.55 | 0.17/0.17 | NBDs + IDs | 1745/2261 | 12; 9/20; 10 |
| UvrA2 | Fpg | 4xATP | 2 | 0.59 | 0.67 | 0.61 | 0.75/0.75 | 0.46 | 0.16/0.17 | NBDs + IDs | 2844/3058 | 20; 16/33; 6 |
| UvrA2 | Fpg | 4xATP | 3 | 0.59 | 0.67 | 0.61 | 0.76/0.76 | 0.56 | 0.15/0.17 | IDs | 1468/1495 | 13; 10/12; 6 |
| UvrA2 | Fpg | 4xATP | 4 | 0.58 | 0.66 | 0.60 | 0.75/0.74 | 0.50 | 0.17/0.17 | NBDs + IDs | 2179/2189 | 39; 12/19; 11 |
| UvrA2 | Fpg | 4xATP | 5 | 0.57 | 0.66 | 0.59 | 0.74/0.73 | 0.54 | 0.15/0.17 | NBDs + IDs | 1491/2335 | 18; 10/24; 17 |
| UvrA2 | EndoIII1 | 4xATP | 1 | 0.60 | 0.68 | 0.62 | 0.76/0.75 | 0.65 | 0.16/0.17 | IDs + Zinc-finger | 2337/1759 | 16; 14/19; 15 |
| UvrA2 | EndoIII1 | 4xATP | 2 | 0.60 | 0.68 | 0.62 | 0.75/0.75 | 0.69 | 0.17/0.16 | IDs | 1095/1100 | 8; 0/10; 5 |
| UvrA2 | EndoIII1 | 4xATP | 3 | 0.59 | 0.67 | 0.61 | 0.74/0.74 | 0.70 | 0.18/0.16 | IDs + Zinc-finger | 1195/1277 | 6; 5/9; 4 |
| UvrA2 | EndoIII1 | 4xATP | 4 | 0.59 | 0.67 | 0.61 | 0.75/0.75 | 0.69 | 0.16/0.17 | IDs + NBD | 967/1977 | 4; 4/19; 13 |
| UvrA2 | EndoIII1 | 4xATP | 5 | 0.59 | 0.67 | 0.61 | 0.74/0.74 | 0.72 | 0.19/0.16 | IDs + Zinc-finger | 1357/1310 | 9; 8/9; 2 |
| UvrA2 | EndoIII2 | 4xATP | 1 | 0.63 | 0.71 | 0.66 | 0.77/0.78 | 0.50 | 0.15/0.18 | IDs | 2299/1770 | 17; 4/19; 4 |
| UvrA2 | EndoIII2 | 4xATP | 2 | 0.63 | 0.71 | 0.66 | 0.78/0.78 | 0.51 | 0.18/0.18 | IDs + Zinc-finger | 1169/1875 | 8; 2/10; 3 |
| UvrA2 | EndoIII2 | 4xATP | 3 | 0.63 | 0.71 | 0.65 | 0.78/0.78 | 0.49 | 0.17/0.15 | IDs | 1831/1716 | 19; 15/14; 10 |
| UvrA2 | EndoIII2 | 4xATP | 4 | 0.62 | 0.70 | 0.65 | 0.77/0.77 | 0.52 | 0.16/0.18 | IDs | 1308/1466 | 8; 1/7; 11 |
| UvrA2 | EndoIII2 | 4xATP | 5 | 0.62 | 0.70 | 0.64 | 0.77/0.77 | 0.51 | 0.18/0.17 | IDs + Zinc-finger | 2269/1624 | 25; 19/15; 7 |
| UvrA2 | EndoIII3 | 4xATP | 1 | 0.58 | 0.67 | 0.61 | 0.75/0.74 | 0.69 | 0.16/0.16 | IDs + Zinc-finger | 2109/2040 | 21; 11/15; 12 |
| UvrA2 | EndoIII3 | 4xATP | 2 | 0.58 | 0.67 | 0.61 | 0.76/0.76 | 0.70 | 0.16/0.16 | IDs + NBD | 2011/2281 | 24; 13/13; 17 |
| UvrA2 | EndoIII3 | 4xATP | 3 | 0.58 | 0.67 | 0.61 | 0.76/0.76 | 0.68 | 0.16/0.15 | IDs | 2073/1767 | 24; 11/19; 8 |
| UvrA2 | EndoIII3 | 4xATP | 4 | 0.58 | 0.67 | 0.61 | 0.75/0.75 | 0.68 | 0.16/0.16 | IDs + Zinc-finger | 1940/2914 | 9; 8/20; 13 |
| UvrA2 | EndoIII3 | 4xATP | 5 | 0.58 | 0.66 | 0.60 | 0.74/0.74 | 0.73 | 0.16/0.15 | IDs + Zinc-finger | 1697/795 | 16; 7/2; 0 |
| UvrA2 | UNG | 4xATP | 1 | 0.61 | 0.69 | 0.63 | 0.76/0.77 | 0.73 | 0.15/0.14 | IDs + Zinc-finger | 3104/2525 | 29; 21/28; 12 |
| UvrA2 | UNG | 4xATP | 2 | 0.61 | 0.69 | 0.63 | 0.76/0.75 | 0.74 | 0.15/0.15 | IDs + NBD | 2193/2361 | 21; 9/21; 4 |
| UvrA2 | UNG | 4xATP | 3 | 0.61 | 0.68 | 0.63 | 0.75/0.76 | 0.73 | 0.14/0.15 | IDs + Zn finger + NBD | 2347/3196 | 31; 5/33; 8 |
| UvrA2 | UNG | 4xATP | 4 | 0.60 | 0.68 | 0.62 | 0.76/0.76 | 0.73 | 0.15/0.15 | IDs + Zn finger + NBD | 3147/2315 | 24; 18/13; 9 |
| UvrA2 | UNG | 4xATP | 5 | 0.59 | 0.67 | 0.62 | 0.75/0.74 | 0.77 | 0.16/0.15 | IDs + NBD | 2170/1552 | 14; 7/9; 4 |

*Nhb: Number of hydrogen bonds. Nsb: Number of salt bridges. Chains 1 and 2 correspond to the two chains of the UvrA dimers. The models exhibiting the highest interacting surface areas (in Å$^2$) and interacting residues are highlighted in green.*

### Table S2: Prediction scores and interface analysis of AlphaFold3 predicted models of UvrA ID domains bound to single domains of BER factors

| UvrA variant | UvrA domain | Partner | Partner region | Rank | Overall ipTM | Overall pTM | Ranking score | Interacting area | Nhb/Nsb |
|---|---|---|---|---|---|---|---|---|---|
| UvrA1 | ID | Fpg | 131-276 | 1 | 0.39 | 0.63 | 0.44 | 800 | 4/2 |
| UvrA1 | ID | Fpg | 131-276 | 2 | 0.27 | 0.56 | 0.37 | 847 | 6/4 |
| UvrA1 | ID | Fpg | 131-276 | 3 | 0.25 | 0.55 | 0.35 | 810 | 6/3 |
| UvrA1 | ID | Fpg | 131-276 | 4 | 0.20 | 0.53 | 0.31 | 787 | 2/5 |
| UvrA1 | ID | Fpg | 131-276 | 5 | 0.21 | 0.53 | 0.31 | 781 | 2/3 |
| UvrA1 | ID | DNA ligase | 321-610 | 1 | 0.28 | 0.61 | 0.37 | 994 | 7/4 |
| UvrA1 | ID | DNA ligase | 321-610 | 2 | 0.27 | 0.60 | 0.36 | 872 | 6/1 |
| UvrA1 | ID | DNA ligase | 321-610 | 3 | 0.25 | 0.60 | 0.35 | 968 | 5/2 |
| UvrA1 | ID | DNA ligase | 321-610 | 4 | 0.25 | 0.61 | 0.34 | 866 | 5/2 |
| UvrA1 | ID | DNA ligase | 321-610 | 5 | 0.24 | 0.60 | 0.33 | 1017 | 2/7 |
| UvrA2 | ID | Fpg | 131-276 | 1 | 0.36 | 0.64 | 0.45 | 687 | 5/0 |
| UvrA2 | ID | Fpg | 131-276 | 2 | 0.29 | 0.60 | 0.39 | 624 | 3/0 |
| UvrA2 | ID | Fpg | 131-276 | 3 | 0.27 | 0.59 | 0.37 | 638 | 4/0 |
| UvrA2 | ID | Fpg | 131-276 | 4 | 0.26 | 0.59 | 0.36 | 634 | 6/0 |
| UvrA2 | ID | Fpg | 131-276 | 5 | 0.25 | 0.58 | 0.35 | 622 | 2/0 |
| UvrA2 | ID | EndoIII1 | 41-162 | 1 | 0.71 | 0.79 | 0.73 | 589 | 5/3 |
| UvrA2 | ID | EndoIII1 | 41-162 | 2 | 0.71 | 0.79 | 0.73 | 575 | 3/0 |
| UvrA2 | ID | EndoIII1 | 41-162 | 3 | 0.71 | 0.78 | 0.72 | 611 | 5/0 |
| UvrA2 | ID | EndoIII1 | 41-162 | 4 | 0.69 | 0.78 | 0.71 | 618 | 6/1 |
| UvrA2 | ID | EndoIII1 | 41-162 | 5 | 0.68 | 0.77 | 0.70 | 588 | 6/2 |
| UvrA2 | ID | EndoIII2 | 34-145 | 1 | 0.49 | 0.71 | 0.53 | 691 | 7/3 |
| UvrA2 | ID | EndoIII2 | 34-145 | 2 | 0.42 | 0.68 | 0.47 | 640 | 4/0 |
| UvrA2 | ID | EndoIII2 | 34-145 | 3 | 0.40 | 0.66 | 0.45 | 690 | 4/0 |
| UvrA2 | ID | EndoIII2 | 34-145 | 4 | 0.25 | 0.60 | 0.32 | 768 | 7/0 |
| UvrA2 | ID | EndoIII2 | 34-145 | 5 | 0.21 | 0.58 | 0.29 | 681 | 7/1 |
| UvrA2 | ID | EndoIII3 | 39-172 | 1 | 0.58 | 0.71 | 0.60 | 703 | 4/2 |
| UvrA2 | ID | EndoIII3 | 39-172 | 2 | 0.48 | 0.67 | 0.52 | 740 | 2/3 |
| UvrA2 | ID | EndoIII3 | 39-172 | 3 | 0.43 | 0.65 | 0.48 | 758 | 6/4 |
| UvrA2 | ID | EndoIII3 | 39-172 | 4 | 0.42 | 0.64 | 0.46 | 774 | 7/2 |
| UvrA2 | ID | EndoIII3 | 39-172 | 5 | 0.38 | 0.62 | 0.43 | 742 | 4/0 |
| UvrA2 | ID | UNG | 2-170 | 1 | 0.50 | 0.68 | 0.55 | 771 | 7/5 |
| UvrA2 | ID | UNG | 2-170 | 2 | 0.48 | 0.67 | 0.54 | 781 | 6/6 |
| UvrA2 | ID | UNG | 2-170 | 3 | 0.42 | 0.64 | 0.49 | 777 | 6/5 |
| UvrA2 | ID | UNG | 2-170 | 4 | 0.17 | 0.54 | 0.25 | 957 | 4/4 |
| UvrA2 | ID | UNG | 2-170 | 5 | 0.15 | 0.52 | 0.24 | 758 | 5/1 |

*Nhb: Number of hydrogen bonds. Nsb: Number of salt bridges.*
